# The Switchmaze: an open-design device for measuring motivation and drive switching in mice

**DOI:** 10.1101/2024.01.31.578188

**Authors:** Clara Hartmann, Ambika Mahajan, Vinicius Borges, Lotte Razenberg, Yves Thönnes, Mahesh M. Karnani

## Abstract

Animals need to switch between motivated behaviours, like drinking, feeding or social interaction, to meet environmental availability, internal needs and more complex ethological needs such as hiding future actions from competitors. Inflexible, repetitive behaviours are a hallmark of many neuropsychiatric disorders. However, how the brain orchestrates switching between the neural mechanisms controlling motivated behaviours, or drives, is unknown. This is partly due to a lack of appropriate measurement systems. We designed an automated extended home-cage, the Switchmaze, using open-source hardware and software. In this study, we use it to establish a behavioural assay of motivational switching in mice. Individual animals access the Switchmaze from the home-cage and choose between entering one of two chambers containing different goal objects or returning to the home-cage. Motivational switching is measured as a ratio of switching between chambers and continuous exploitation of one chamber. Behavioural transition analysis is used to further dissect altered motivational switching. As proof-of-concept, we show environmental manipulation, and targeted brain manipulation experiments which altered motivational switching without effect on traditional behavioural parameters. Chemogenetic inhibition of the prefrontal-hypothalamic axis increased the rate of motivation switching, highlighting the involvement of this pathway in drive switching. This work demonstrates the utility of open-design in understanding animal behaviour and its neural correlates.

## Introduction

Motivated behaviours, like feeding, drinking and social interaction are stereotypical behaviours critical for survival^1,2^. Motivated behaviours are generated by neural mechanisms, termed drives^3,4^. We note that this may be distinct from other definitions, such as drive as a variable that is reduced during a behaviour^5^. In health, different drives alternate to meet internal needs, while also being controlled by the availability of goal objects^6^, conscious effort and complex needs, like hiding actions from competitors^7^. High and low drive switching at a rapid timescale can be adaptive in different circumstances. Frequent switching is ideal when goal objects are presented in unexpected locations, such as when exploring a new or altered area, whereas infrequent switching should be preferred when exploiting a familiar environment during a high need state^7,8^. Inability to switch drives rapidly could account for the behavioural rigidity seen in many neuropsychiatric disorders, and the slowness of behavioural completion in neurodegenerative disorders^9^. In order to study drive switching, we set out to measure switching between motivated behaviours in mice, i.e., motivational switching.

Behavioural flexibility is typically studied in mice using approach/avoid decisions^10,11^ or simplified set-shifting tasks under water deprivation and after arduous training regimens^12,13^. However, the fragmented nature of mouse behaviour complicates translational use of cognitive flexibility tasks like set-shifting^12–14^. We sought to improve on this by analysing flexibility of spontaneous motivated behaviours, feeding, drinking and socializing. Most previous studies of behavioural flexibility rely on teaching rodents a motor action such as lever pressing that is unlikely to be encountered in the wild. Instead, navigating a familiar walled environment is natural for mice^15,16^. Therefore, we set up a naturalistic sequential foraging task which discretizes motivational switching – the Switchmaze^17^. We quantify motivational switching with a behavioural metric derived from the ratio of single probe entries to continuous exploitation runs, termed *motivation switching rate (MSR)*. Here, we demonstrate the utility of this approach by showing that it can reveal new information about the structure of natural motivational switching and its neural control.

There is a high degree of commitment in behavioural cycles^2^. That is, once a motivated behaviour (such as feeding) is initiated, its termination likelihood is initially low and increases thereafter^18^. This is also reflected in motivated behaviours being resistant to distractor stimuli^19^, and need-based meal length (rather than meal frequency) regulation^20^. We analyse the degree of commitment using run lengths (how many behavioural cycles until a run of one motivation is terminated) and likelihoods of motivational transitions.

Hypothalamic neurons are critical components of the drives for feeding, drinking and social behaviours^21^. However, how the brain coordinates switching between drives is a major unknown. Furthermore, most studies use overly simplified, shoe-box sized environments with highly divergent goal object availability from that encountered in natural habitats. E.g., food availability is often constant and unlimited in most recordings and throughout the animal’s life. As a result, naturalistic drive switching is highly understudied.

Drive switching likely involves the prefrontal cortex (PFC), which is involved in sequencing goal-directed actions and is densely connected with the hypothalamus^22,23^. We monitored motivation switching in discretized, repeatable behavioural cycles, while chemogenetically inhibiting the hypothalamus or PFC projections to the hypothalamus, in order to investigate their contributions to drive switching. Our results suggest that MSR is a useful parameter, arising particularly clearly from the Switchmaze, and that PFC→hypothalamus neurons are part of a drive network regulating motivation switching by promoting continuous feeding bouts.

## Materials and methods

### Switchmaze

Build instructions for the Switchmaze, a bill of materials and code are available in the Supporting Files. The apparatus (Figure 1) discretizes behavioural cycles of feeding and drinking, which animals perform one at a time in a foraging environment separated from the home-cage by a single-entry module. Once inside the foraging environment, upon each entry to a goal area (a trial), food and drink were only available in a ‘quantum’ of either one 14 mg pellet (Bio-Serv™ Dustless Precision Pellets™ for Rodents, F05684) or one ~10 μl drop of water. After goal entry, the animal has one ‘return’ path available to the start position, which differs from the entry path. Once the animal is in the start position, the availability resets for both food and drink, and the animal also has the option to return to the home-cage. Thus, switching between three fundamental motivated behaviours can be followed when the animal is in the start position. Mice live in the apparatus for several days on a 12/12 light dark cycle (lights off at 9 am). The Switchmaze records entries to the foraging environment, start point, and feeding and drinking areas, as well as the consummatory actions, pellet retrieval, drinking and running wheel use. Health and welfare monitoring and maze cleaning was done at least once daily and safe operation was monitored on an overhead camera. The apparatus was in an isolated procedure room for all recordings.

**Figure 1.**
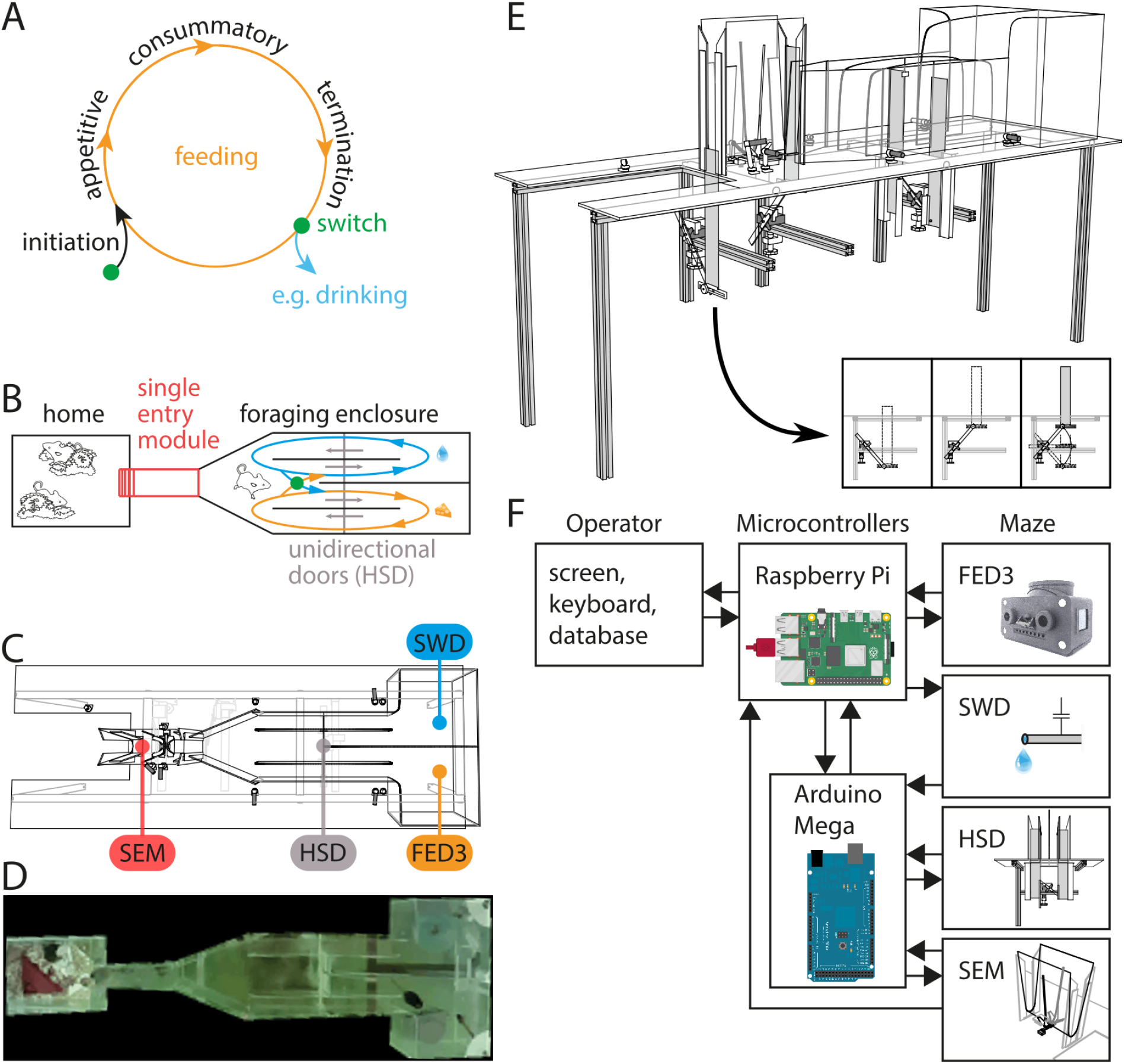
Designing the Switchmaze to measure behavioural cycles spatially. A, Concept of feeding as a behavioural cycle, following^1,2^; B, Schematic of feeding and drinking cycles separated spatially in the Switchmaze; C, CAD model from above; D, Picture of maze during operation with a mouse retrieving a food pellet (bottom right); E, CAD model from side; inset shows how the upward closing doors operate; F, Wiring diagram of electrical components. FED3, Feeding Experimentation Device 3^27^; SWD, Sensing Water Dispenser^28^; HSD, Horizontal Sliding Door; SEM, Single Entry Module^29^.

### Stereotaxic surgery

30 wild-type C57BL6 male mice were used in this study. All experimental procedures were approved by the Netherlands Central Committee for Animal Experiments and the Animal Ethical Care Committee of the Vrije Universiteit Amsterdam (AVD11200202114477). Of the 30 mice, 9 were controls (ctrl), 10 expressed inhibitory designer receptors exclusively activated by designer drugs (DREADDs) in the hypothalamus (H-hM4Di), and 11 expressed inhibitory DREADDs in PFC→hypothalamus projection neurons (PFC-hM4Di). Mice were anesthetized with an intraperitoneal (i.p.) injection of a mixture of fentanyl (0.05 mg/kg), medetomidine (0.5 mg/kg) and midazolam (5 mg/kg) in saline, the scalp was injected subcutaneously with lidocaine, opened, and 0.2 mm craniotomies were drilled bilaterally at 0.9 mm lateral, 1.4 mm posterior from Bregma. For medial PFC injections of AAV8-syn-DIO-hM4Di-mCitrine, additional craniotomies were drilled bilaterally at 0.4 mm lateral, 1.8 mm anterior from Bregma, and 0.4 mm lateral, 2.3 mm anterior from Bregma. A pulled glass injection needle was used to inject the below doses of virus at a rate of 10-50 nl/min.

All H-hM4Di and PFC-hM4Di animals received hypothalamic injections bilaterally 5.4 mm deep in the brain. Out of the 9 H-hM4Di animals, 4 were injected with AAV9-CMV-Cre-tdTomato (10^12^ GC/ml) and AAV8-hSyn-DIO-HA-hM4Di-mCitrine (10^13^ GC/ml) (mixing ratio: 1:5; injection volume: 30-150nl). The remaining 5 animals were injected with AAV8-hSyn-hM4Di-mCherry (2*10^13^ GC/ml; injection volume: 30 nl). As there were two groups of H-hM4Di being pooled, we checked if the effect in Figure 6A was different between these groups, and found that it was not (the MSR change due to C21 in the Cre:DIO-hM4Di mixture group was −9.4 ± 49.5 %, and in the hSyn-hM4Di group it was −5.6 ± 42.5 %, P=0.9).

All PFC-hM4Di animals, received hypothalamic injections of AAVrg-hSyn-Cre-P2A-tdTomato (1.5*10^13^ GC/ml, injection volume: 30nl) as well as bilateral medial PFC (mPFC) injections of AAV8-hSyn-DIO-HA-hM4Di-mCitrine (10^13^ GC/ml). The mPFC injections were split into three locations per hemisphere: Two at 1.8 mm anterior from Bregma, 0.4 mm lateral at depths 2.0 mm and 1.5 mm (150 nl), and one dose at 2.3 mm anterior from Bregma, 0.4 mm lateral at depth 1.75 mm (injection volume: 150 nl at each location).

Injection needles were kept in place for 20 min in hypothalamus and 3-5 min in mPFC before withdrawing. After the injections, an RFID chip (Sparkfun SEN-09416) was implanted under the chest skin, the wounds were closed with tissue glue, anaesthesia was antagonized with an injection of flumazenil (0.1 mg/ml) and atipamezole (5 mg/ml) in saline (i.p.). Animals received 0.05 mg/ml carprofen in drinking water for 2-4 days as post-operative pain medication.

### DREADD manipulation experiments

Animals were injected with saline on the first experiment day and 5 mg/kg C21 (agonist of hM4Di) dissolved in saline on the second day. Behavioural parameters were measured over a 6-h window as the effect of C21 is likely to last at least that long^24,25^, and MSR can be highly variable in short time windows which may contain a low amount of trials.

### Histology

We examined viral targeting with post-experiment histology (see Supplemental information for micrographs of transgene expression in target areas) in 70 μm coronal sections cut from immersion-fixed brains extracted into 4% PFA after an overdose of pentobarbital solution administered under isoflurane anaesthesia. In PFC-hM4Di specimens, the hypothalamic injection centroid was always within the lateral hypothalamic area (LH) with a minority of cases with expression elsewhere (1/11) or difficult to detect expression (1/11). In all cases the mPFC layer 5 (L5) neurons expressed hM4Di-mCitrine, with difficult to detect expression in 1 case out of 11 (same individual as the above difficult to detect hypothalamic expression) and abundant expression in all others. All animals were included in the analysis. In H-hM4Di specimens, the injection centroids were always within the LH with a minority of cases with stronger expression in one hemisphere (2/9). As all cases contained hM4Di-expression in the target regions, we included all cases in analysis.

### Statistics

Of the 30 animals in this study, 4 were in the representative dataset in Figures 2A-E,G,H and 3; 13 in the representative dataset in Figure2F,I; 20 in the habituation dataset in Figure 4 (left plots); 22 in the environmental challenge dataset in Figure 4 (middle and right plots) and Figure 5, where 15 animals were the same as in the habituation dataset (Figure 4, left plots); 30 in Figure 6; 11 in Figures 7 and 8. Data were analyzed using Matlab R2019a and reported as mean ± SD. For the chemogenetic experiments we used two-way repeated measures ANOVA with time (vehicle or C21) as the within-subjects factor, and cohort (ctrl, H-hM4Di or PFC-hM4Di) as the between-subjects factor. This was done by fitting a ‘WithinDesign’ repeated measures model (fitrm) in Matlab with Cohort as the predictor variable, followed by repeated measures analysis of variance (ranova). When a significant cohort-time interaction was found, three follow-up paired t-tests were used with Bonferroni-corrected significance threshold of 0.0167. Paired t-tests (Bonferroni-corrected significance thresholds: Figure 4, 0.01; Figures 5 and 7, 0.0056) and Wilcoxon rank sum tests were performed in Matlab. In Figure 4, a 4-hour time bin was used as it was estimated to be the lowest time resolution that provides reasonably precise values of MSR. As a ratio between singles and runs, in blocks that occur somewhat stochastically across hours (see Figure 2A), this parameter becomes highly variable at short time bins as the absolute values in numerator and denominator may vary.

**Figure 2.**
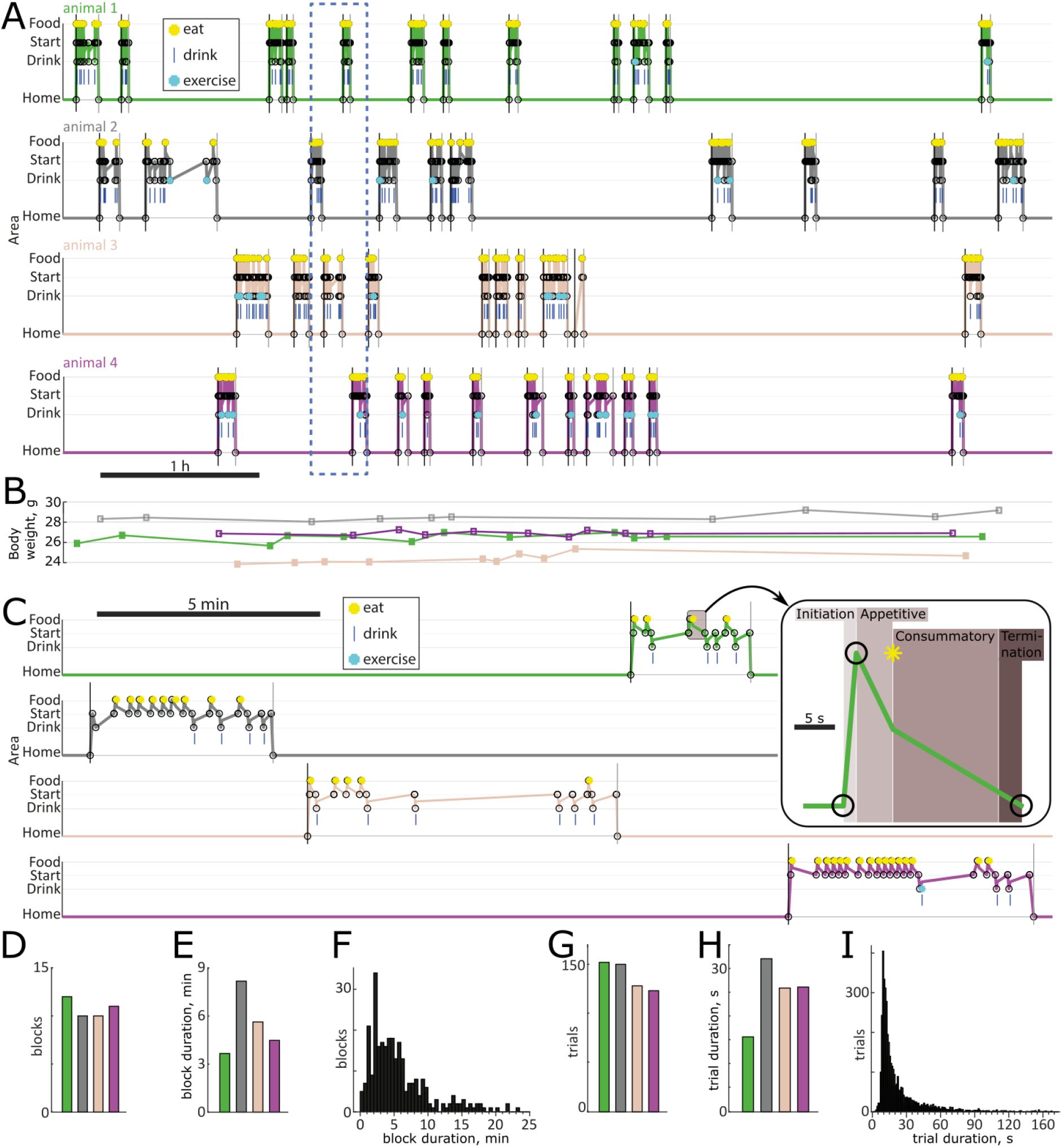
Spontaneous behaviour of four mice habituated to live in the Switchmaze. A, Ethograms of four mice entering the foraging environment one at a time for an open-ended block. B, Weights of the mice were recorded at each entry to the foraging area. C, One block for each animal shown in detail in the expanded time window (dashed box in A). Inset shows one trial from animal 1 at a further expanded time scale to highlight the fractionation of the behavioural cycle (as in Figure 1A). D,E,G,H, basic metrics for the 6 h session shown in A, colour coded by mouse. D, number of blocks; E, mean block duration; F, block durations across 13 animals in a 24 h period. G, number of trials; H, mean trial duration; I, trial durations across 13 animals in a 24 h period.

**Figure 3.**
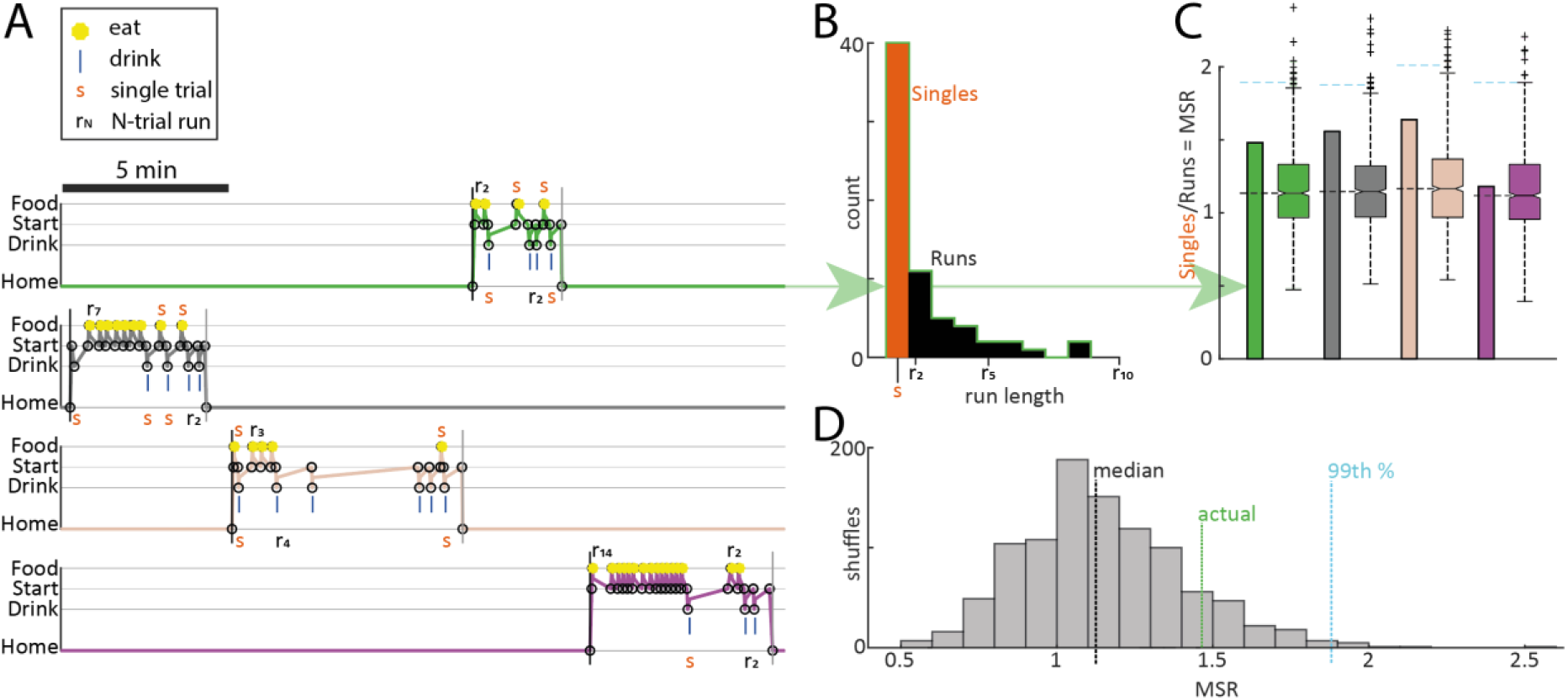
Motivational switching. A, Ethograms for four mice entering the foraging environment one at a time for an open-ended block. ‘Singles’ and runs labelled for clarity. B, For animal 1 only, distribution of singles and runs by length. C, MSR for each animal (bars, number of singles divided by number of runs of any length) and MSR expected by random chance (box plots are MSR from 1000 random permuted behaviour sequences for each animal; dashed horizontal lines denote median (black) and 99th percentile (cyan)). D, Distribution of MSRs calculated from shuffled behaviour sequences for animal 1.

**Figure 4.**
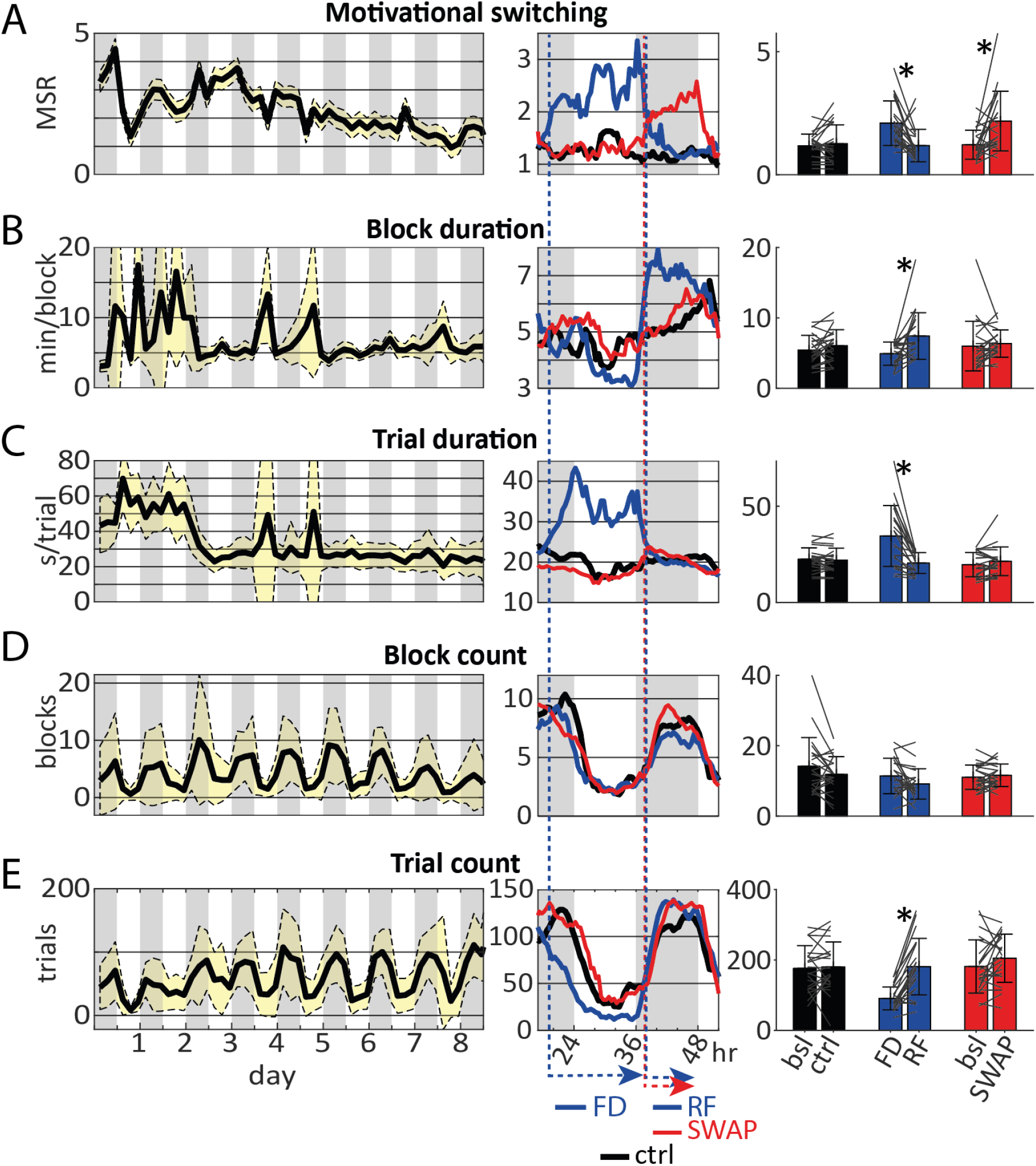
Stability and dynamics of Switchmaze parameters during habituation and challenges. A, MSR; B, Block duration; C, Trial duration; D, Block count; E, Trial count. Time series data (left and middle panels) have a 4 h bin width and bar graphs (right panels) are measured from the last 6 h before lights-on. Dark phases of the light-dark cycle are marked in grey. N=20 for habituation plots (left), N=22 for the challenges (middle and right; 15 animals were in both datasets). FD, food deprivation; RF, re-feeding; SWAP, swap goals. *, p<0.004.

**Figure 5.**
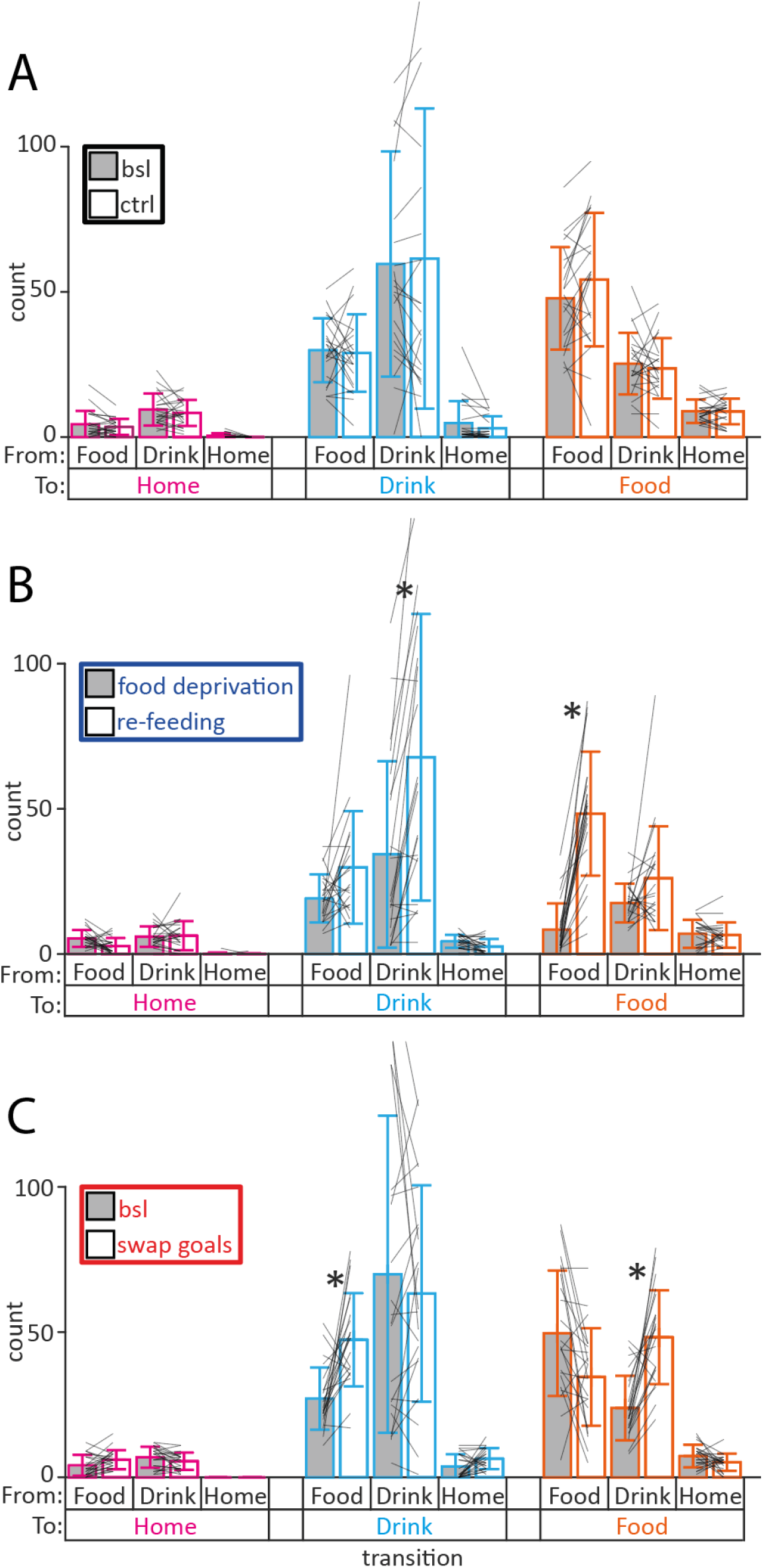
Trial transitions during 6 h before lights-on. Same raw data as Figure 4 right side plots. A, Baseline and control; B, Food deprivation and re-feeding; C, Baseline and swapped goals. * p<0.0056

**Figure 6.**
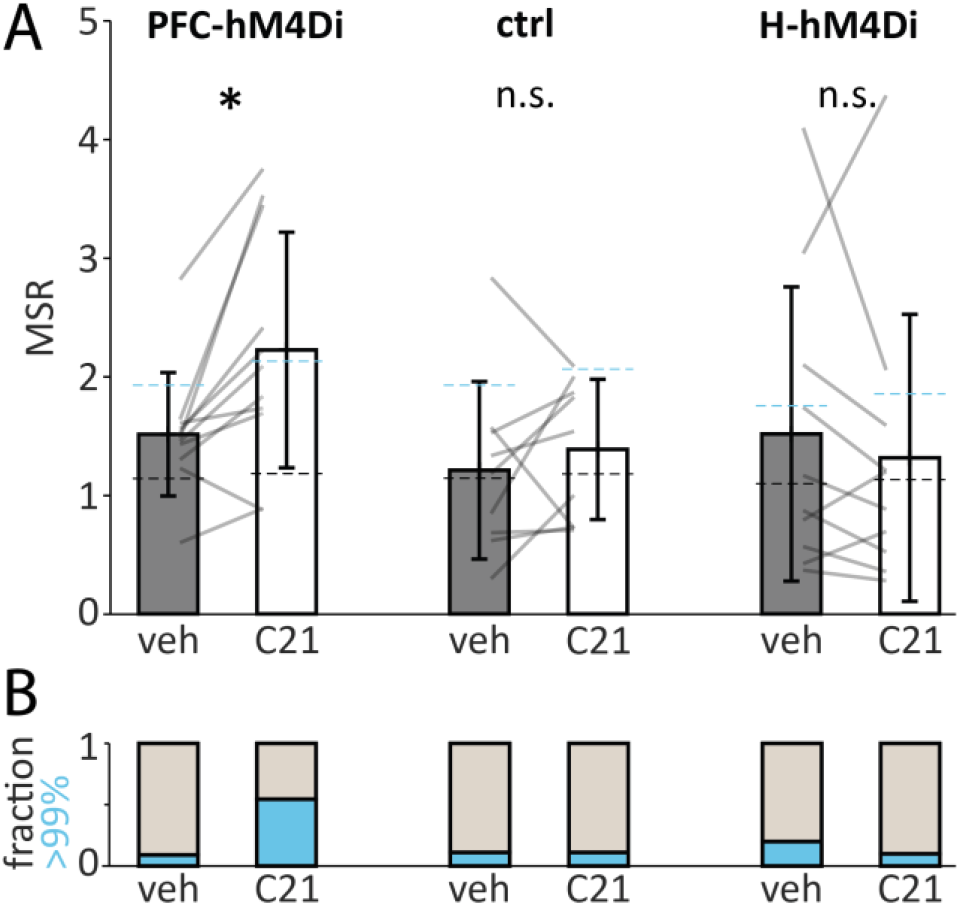
Chemogenetic modulation of motivational switching. A, MSR quantified as singles/runs in PFC-hM4Di (left, n=11), ctrl (middle, n=9) and H-hM4Di cohorts (right, n=10) during the 6 h after vehicle or C21 injection. Two-way repeated measures ANOVA revealed a significant cohort-time interaction F(2, 27)=3.99, p=0.03 and paired t-tests revealed the effect was in the PFC-hM4Di cohort, * p=0.007. Dashed lines denote chance level arising from the mean of medians (black dashed line) and mean of 99th percentiles (cyan dashed line) of 1000 random-permuted behavioural sequences for each animal. B, Fraction of animals with switching rate higher than the 99th percentile of its random permutations in cyan.

**Figure 7.**
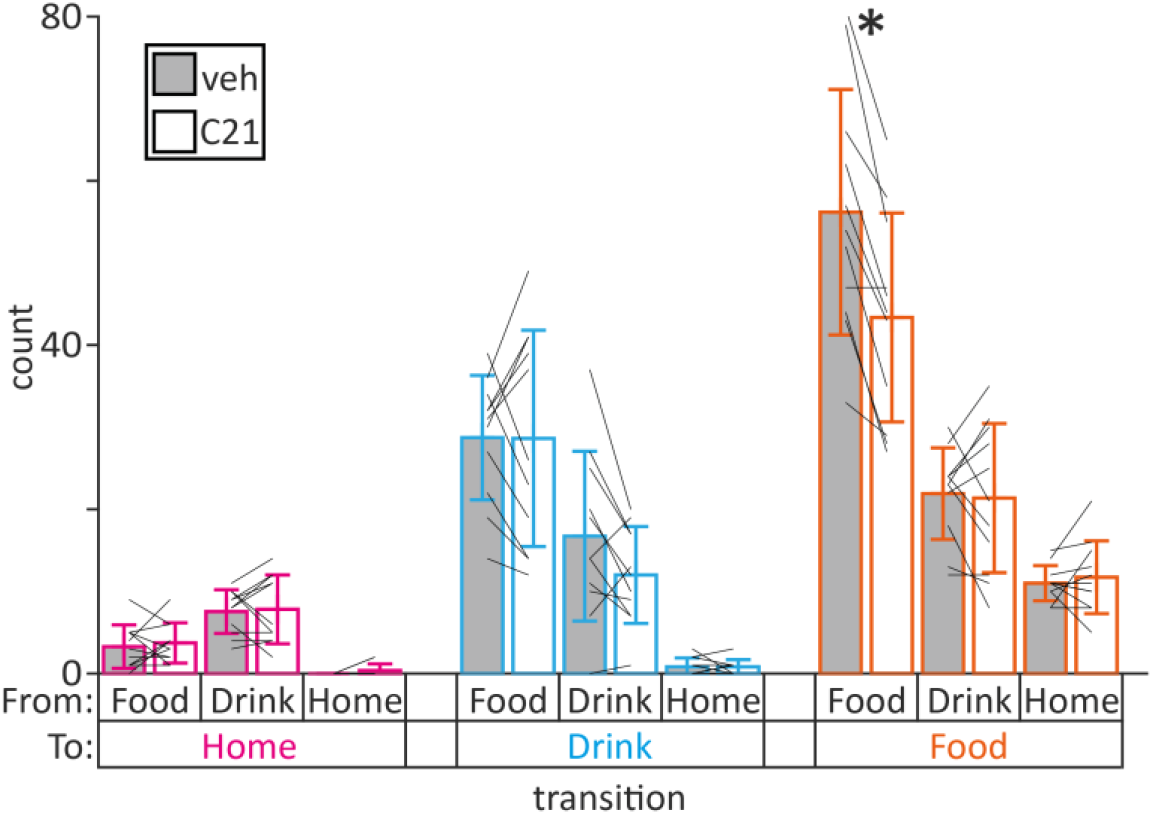
Food-to-food trial transitions are decreased by PFC inhibition. Same raw data as Figure 6 PFC-hM4Di panels. * p=0.00009

**Figure 8.**
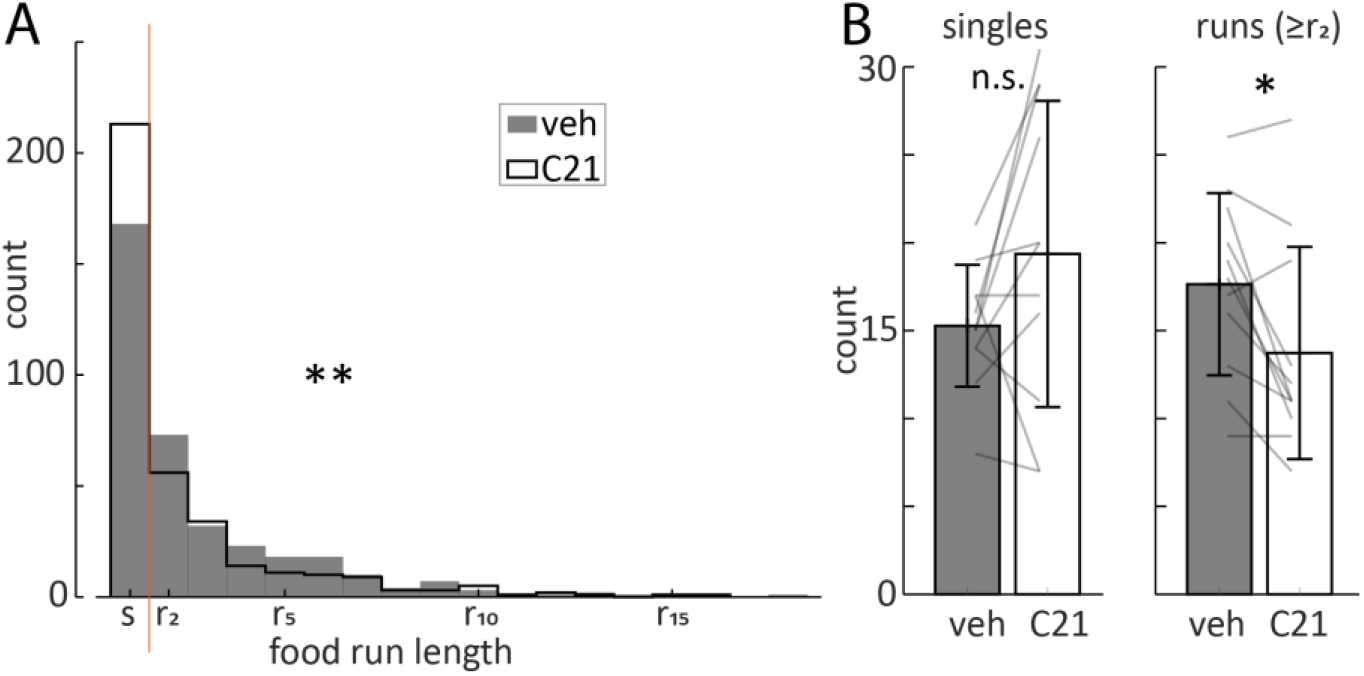
Food runs are decreased by PFC inhibition. A, Run length distribution for food runs in the PFC-hM4Di cohort (n=11) from the same raw data as Figure 6 PFC-hM4Di panels. B, Food singles and runs (more than 1 cycle) analysed for each animal. ** p=0.002, * p=0.01

### Data, script and code availability, and supplementary information

Data, code, and supplementary information are available at https://doi.org/10.17605/OSF.IO/YB67Q.

## Results

### Designing the Switchmaze

Feeding, drinking and social interaction are cyclical behaviours that can be divided into initiation, appetitive, consummatory and termination epochs^1,2^ followed by a potential switch to another behavioural cycle (Figure 1A). In order to isolate these phases, in particular the switch epoch, we designed a foraging area that spatially separates feeding and drinking cycle phases (Figure 1B). To enforce directionality of the spatial cycles, we used unidirectional doors (termed a horizontal sliding door unit, HSD) that operate based on beam-break detector signals that determine the mouse’s location in the maze. Thus there was one entry and one exit pathway to the food and drink areas, which is similar to natural burrows of mice^15^. On each entry, the mouse could access one quantum of water (10 μl) or food (14 mg). These were selected in order to get multiple consecutive entries to each area for completing a typical *ad libitum* consumption bout measured in laboratory conditions (60 μl water^26^, 220 mg food^27^), to mimic estimates of wild mouse feeding patterns (approximately 200 small meals each day^16^), and to induce approximately the same amount of entries into both areas, such that the likelihood of switching at each re-entry to the start position would be approximately equal. After retrieving food or water, the mouse has one path available to the start position, where availability resets and access to the home area is always available.

To allow detection of individual mice, each mouse was tagged with a radio frequency identification (RFID) tag. To pass only one mouse into the foraging enclosure only when it was unoccupied, a single entry module (SEM) was designed^29^. The SEM uses three types of sensors: an RFID antenna, weight sensor, and beam-break sensors. These allow for identifying a single animal in a safe position within the module, such that the upward closing doors (doors that move downward below the floor plate to open the passage, and upward to block the passage; Figure 1E inset) can close without risk of lifting a mouse on top of them. As there is no ceiling, there is also no danger to animals from the upward closing doors, and the device is, in principle, compatible with tethered recordings (Figure 1E).

While one animal was inside the foraging enclosure, the others stay in the home area, which was a standard Perspex home-cage bottom with bedding and several shelters. The maze was built mostly from transparent acrylic sheet in order to allow flexible placement of beam break detectors and sequential assembly with chemical welding (Figure 1C-E). Other components included 3D printed PLA components, aluminium rail and sheet, and off-the-shelf electronics (for full details and building instructions, see the Supplement). The overall idea was to mimic natural mouse burrows and cavity walls of human dwellings.

Food was provided with a FED3 pellet dispenser^27,30^, and water with a custom lick-activated water dispenser^28^ (Figure 1D,F). Doors were operated with servo motors controlled by an Arduino microcontroller, and master control and data logging were done on a Raspberry Pi computer using custom Python code (Figure 1F). The Arduino was also used to measure animal location based on beam break detectors and to measure licking contact at the water spout through capacitive sensing. Thus, tasks were divided between the Raspberry Pi and Arduino to make use of their strengths, i.e., logging data and interfacing with the operator (Raspberry Pi), and controlling servo motors and recording analogue signals (Arduino).

### Measuring basic metrics in the Switchmaze

We started by observing the behaviour of C57BL6 mice housed in the Switchmaze for up to a month in cohorts of 2-4 animals. For the first two days, water was available *ad libitum* in the home-cage and 2g/animal of dry chow was provided on the home-cage floor. This was done to ensure adequate food and water intake during the time when entering the foraging environment is a new action. Therefore, mice would first enter the foraging environment due to exploration. In 6-48 h, they learned to obtain food and water from the maze. During baseline operation, an animal consumed on average 189 ± 49 food pellets (14 mg) in a 24 h period, which corresponds to the typical number of small meals wild mice are estimated to eat in a night^16^. Upon most entries into the food pod, a pellet was consumed (93.9 ± 10.1 % of entries). 210 ± 130 water drops (10 μl) were retrieved and drinking occurred upon 98.5 ± 2.3 % of entries into the water pod. In a 24 h period, an animal ran on the running wheel on average 6.7 ± 13.3 revolutions during 5 ± 7 entries into the water pod which accounted for 4.0 ± 6.5 % of the entries. Therefore, despite the appeal of running wheels even to wild mice^31^, use of the running wheel in the Switchmaze was marginal and was excluded in further analyses. Entries to the food pod (201 ± 49) and water pod (212 ± 130) were approximately balanced over a 24 h period.

After habituation (>7 days), animals exhibited stereotyped entries into the foraging environment (Figure 2A) to eat and drink, followed by exit back into the home-cage. We termed these foraging area visits *blocks*. Animals were automatically weighed at the start of each block (Figure 2). Blocks lasted on average 6.3 ± 8.0 min (Figure 2C-F) and tended to occur in clusters across the group, such that the animals spent time together in the home-cage (Figure 2A). This suggests that the animals returned to the home-cage due to a strong social motivation. They would rarely stay in the foraging environment for longer than 17 minutes (Figure 2F; 95^th^ percentile = 16.9 min).

During each block, an animal typically made multiple entries to the food and drink areas (Figure 2C), termed *trials*. Trials lasted on average 26.3 ± 34.1 s (measured from start position to return to start position, i.e., a full behavioural cycle through one goal area, Figure 2C inset), and rarely more than 71 s (Figure 2G-I; 95^th^ percentile = 70.7 s). These short cycle durations, together with the brief blocks, suggest that the foraging behaviour was economical and purposeful.

### Measuring motivational switching

To satisfy a need such as hunger, an animal would be expected to enter the food area repeatedly, as only one quantum of consumption was available at each trial. These repeated entries, termed runs, occurred to both food and drink areas (Figure 3A). However, the most common run length was one trial, which can be seen as a behavioural phenomenon controlled neurally by *switching the underlying drive* before and after the trial. Similar single trials are common in rodent behaviour, in particular during trained performance of simple tasks by highly skilled animals^8,13,32^. This seemingly counterproductive behavioural stochasticity may be part of a behavioural camouflage mechanism evolved to elude competitors^7^. In order to capture the prevalence of these ‘singles’, we analysed the ratio of singles to runs (from here, we define a run as a sequence of repeated food or drink trials longer than one). This motivation switching rate (MSR) was on average 1.4 ± 0.9 (Figure 3B,C) after habituation (>7 days). To test if motivational switching was different from random, we randomly shuffled the trial sequence and re-calculated MSR 1000 times for each animal (Figure 3C,D). The arising distribution encompassed the actual MSR in most cases (only 4/30 animals had MSR higher than the 99^th^ percentile of the shuffled data). This suggests that spontaneous motivation switching is optimized to appear random, which could conceivably function to decrease the predictive information available to competitors and predators.

### Habituation and environmental control of motivational switching

We next asked how motivational switching changes when the mice habituate to the Switchmaze. During the first eight days, MSR was high in the beginning and then settled toward a steady state value around 1.4 (Figure 4A). This further suggests that while high switching rates above the random regime may be useful for exploration of a new environment, the animals optimize motivational switching toward the median of the random distribution (Figure 3D). Durations of blocks and trials also settled to a steady baseline during the first week (Figure 4B,C) and their frequencies settled into a steady diurnal pattern (Figure 4D,E). Blocks may be thought of as meals, because their overall number during the light and dark cycle matched previous estimates of meals in mice with *ad libitum* access to a FED3 pellet dispenser^27^.

After habituation, we tested how environmental challenges affect the stabilized metrics. Removing the food dispenser (Figure 4; food deprivation, FD) rapidly increased MSR and trial duration, while decreasing trial number, reflecting increased food seeking exploration. These parameters returned to baseline when the food dispenser was returned, 20 hours after removal (Figure 4; re-feeding, RF). Additionally, block duration increased, as more time was spent in the foraging environment in homeostatic re-feeding.

We simulated an environmental uncertainty, similar to depleting resources in patch foraging, by swapping the food and drink areas (Figure 4; SWAP). This resulted in increased MSR without other parameter changes, demonstrating that the MSR is a useful indicator of behavioural structure changes that would not be evident from traditional metrics.

As the increased MSR could arise from increased transitions between drives or decreased continuous runs of one drive, or both, we analysed the transition counts for the environmental challenges (Figure 5). In control conditions transition counts remained unchanged between the baseline and experimental day (Figure 5A). Re-feeding increased both food-to-food and drink-to-drink transitions (Figure 5B), as expected from homeostatic re-feeding^33^. Swapping goals increased food-to-drink and drink-to-food transitions, while food-to-food and drink-to-drink transitions remained unchanged (Figure 5C), suggesting that after entering the intended goal area the animals could re-locate it. These data demonstrate the utility of identifying behavioural sequence changes through MSR, and dissecting them further with transition analysis.

### Neural control of MSR, drive switching

To assess the potential of the Switchmaze for identifying neural underpinnings of motivational switching, we performed two chemogenetic loss-of-function experiments on neural circuits known to be involved in feeding in a non-trivial way. A broad perifornical region of the hypothalamus, containing the lateral, dorsomedial and tuberal areas, regulates feeding, potentially through primary effects on arousal, locomotion and metabolism^34–36^, and contains intermingled feeding promoting and inhibiting neural populations^37–39^. The medial PFC sends axonal projections to this hypothalamic area, affecting feeding in a complex manner depending on behavioural context^40,41^. To test the role of these neural populations in drive switching, we expressed the inhibitory DREADD, hM4Di in either PFC output neurons to the hypothalamus (PFC-hM4Di) or in the perifornical hypothalamus (H-hM4Di). As a control cohort, we used wild-type mice that did not express a transgene, and which were interleaved in groups of hM4Di expressing mice that lived in the Switchmaze.

Basic behavioural metrics were not changed in PFC-hM4Di or H-hM4Di mice upon activation of the inhibitory DREADDs with C21, as two-way repeated measures ANOVA tests showed no significant cohort-time interaction for food consumed (F(2,27)=0.61, p=0.55), block count (F(2,27)=0.66, p=0.52), trial count (F(2,27)=0.50, p=0.61), block duration (F(2,27)=0.75, p=0.48) or trial duration (F(2,27)=0.69, p=0.51). However, MSR was altered significantly (cohort-time interaction F(2,27)=3.99, p=0.03) and paired t-tests revealed the effect was a 46.5 ± 45.5% increase in the PFC-hM4Di cohort (p=0.007, Figure 6A). The average switch rate exceeded the 99^th^ percentile of switch rates expected from randomly permuted trial sequences (cyan dashed lines in Figure 6A). This coincided with 55% (6 out of 11) of PFC-hHM4Di animals increasing their switch rate above the 99^th^ percentile of their distribution of random sequence switch rates (Figure 6B). This result supports previous findings suggesting that a behavioural role of the PFC is to regulate decision sequence stochasticity^7^.

To find out if the change in switching rate of PFC-hM4Di animals was due to decreased repetitive cycles, increased transitions, or both, we analysed their trial transitions. The only significantly changed transition was Food-to-Food, which was decreased by 22.7 ± 11.5% (p=0.0009, Figure 7). This suggests that PFC→hypothalamus projections promote repetitive feeding.

To test if this decrease in food-to-food transitions reflected shortened feeding runs or less of them or both, we analysed the structure of food runs in vehicle and C21. The mean length of runs to the food area did not change (+2.4 ± 14.9%, p=0.6), nor did the mean number of single entries to the food area (+26.2 ± 49.3%, p=0.1). However, the mean number of food runs decreased by 21.9 ± 21.5% (p=0.01, Figure 8). These results suggest that maintenance of a proportion of food runs is controlled by PFC→hypothalamus projection neurons.

Together, these data suggest that PFC→ hypothalamus projection neurons selectively promote food runs rather than single feeding cycles, and this effect maintains MSR in the random regime.

## Discussion

The Switchmaze is a low-cost (about 1100 EUR), open source, ethologically relevant semi-natural environment. Its modular design allows for changing and modifying the affordances to enable diverse experiments, and the simple construction techniques can be readily used to create new goal modules. Mice can stay in the apparatus indefinitely, as all the affordances they have in typical home-cages are provided, and the maze is a highly enriched environment. Such liveable experimental environments are ideal for probing internal models^42^ and schemas^43^ used in daily life and have great potential for generating new rodent models of disorders where a complex daily environment is thought to affect progression, e.g., obesity and eating disorders. The Switchmaze discretizes daily survival behaviours for convenient measurement by turning the behavioural cycles into cyclic routes that mimic foraging paths. This is in stark contrast to typical behavioural experiments where the animal is tested for minutes to hours in a small enclosure with an endless supply of an affordance, followed by return to a different home-cage environment. We aimed to make the device compatible with tethered recordings by using upward closing doors and not including a ceiling. However, the length of the maze may pose additional challenges for the use of tethered recordings, and users interested in this may wish to shorten the layout of their build, or use counterweighted tethers^44^.

Users interested in building a Switchmaze should be aware of similar designs by other groups^27,30,45–54^. The most relevant for projects like Switchmaze are the RFID enabled Autonomouse^45^, Eco-HAB^46^, and several devices from the Murphy lab^47–50^; the FED3 pellet dispenser^27,30^; and the modular open-source solutions Autopilot^51^, pyControl^52^ and LabNet^53^. Of note, automated behavioural devices greatly reduce markers of stress^55^, and enable recordings in the wild^31,56^. Currently, more than 250 similar projects are listed at https://edspace.american.edu/openbehavior/.

We have demonstrated a behavioural measurement of motivational switching in an ethological semi-natural setting. As the switching rate index, we selected a metric that captures the probe trials (singles) which are stereotypical in rodent behaviour^8,13,32^. This metric is highly useful as it provides information that is otherwise missed by traditional metrics (Figure 4, SWAP condition). By observing distributions of random shuffled motivational sequences (Figure 3D), we found that mice optimize their MSR to near the median where the highest number of sequences would be found by chance. This behavioural pattern may be useful to hide their motivations from other animals. As this reduces their predictability, which could be exploited by competitors, it may be a form of behavioural camouflage. This could be tested further by correlating MSR with social rank and testing it in the presence and absence of predator cues. As social factors, like hierarchy, were not controlled for, they may be a potential confound of our results.

During habituation to the maze, i.e., exploration of a novel environment, MSRs started high (predictable) and gradually came closer to the median of the random distribution (Figure 4A). During the life-threatening challenge of food deprivation, switching rates rose (predictable) and fell again during re-feeding, suggesting that behavioural camouflage through optimized drive switching (BeCODS) can be discarded when necessary and rapidly reinstated (Figure 4A). An intriguing possibility, consistent with a lower bound of MSR (Figure 4A), is that an independent drive switching mechanism pulls motivational switching toward the median of a random regime, while other independent mechanisms can elevate it in response to need (FD) or environmental instability (SWAP and habituation). The potential survival benefits from BeCODS likely depend on a stable territory, as goal swapping increased MSR. Chemogenetic inhibition of PFC→hypothalamus projection neurons increased MSR above the random regime (Figure 6). As the neocortex evolved relatively recently, this suggests that BeCODS may be an evolutionarily recent refinement of ancient hypothalamic networks.

The increased switching rates due to PFC-hM4Di were accompanied by decreased food-to-food transitions (Figure 7), similar to food deprivation (Figure 5B), suggesting that PFC-hM4Di affected maintenance of the feeding drive. Of note, we did not find an overall change in food consumed, similar to a recent study of chow consumption in the home-cage^41^. The pattern of motivational transitions affected by PFC-hM4Di was distinct from that induced by swapping the goal modules (Figure 5C), which was accompanied by increased food-to-drink and drink-to-food transitions, as expected from an action-outcome mismatch. As such a mismatch would be expected to arise from amnesia, it appears likely that PFC-hM4Di did not reduce memory function. This is in line with a recent study showing improved go/no-go task performance during inhibition of PFC→LH projections^57^. In summary, the motivational pattern changed in a way that suggests a role for PFC layer 5 → hypothalamus connections in sustaining the feeding drive and, through this effect, decreasing the predictability of the animal’s behavioural sequences.

## Acknowledgements

We thank Tinco Brouwer and Julian van der Velde at the Vrije Universiteit Amsterdam mechanical and electronic engineering workshops for technical assistance, as well as Alexxai Kravitz and co-workers for developing the FED3 pellet dispenser^27,30^, and SparkFun Electronics for developing the open source RFID and weight sensors which greatly facilitated the development of the Switchmaze. We thank Jesse Jackson, Denis Burdakov, Rebecca Jordan, Andre Maia Chagas and Marion Rivalan for useful comments.

