## Supplemental Information for "The Switchmaze: an open-design device for measuring motivation and drive switching in mice"

#### List of supplemental files:

**Build instructions (in this file)**
**CAD models for viewing overall build**
**Code**
**Data and analysis scripts**
**Histology (in this file)**
**Part files for 3D-printing**
**Part files for laser cutting**

### Build instructions for the Switchmaze

The Switchmaze consists of a home cage coupled through a single entry module to a foraging environment where an animal can serially retrieve a quantum of food or water from an isometrically placed decision point (Figure 1). It can be built by an advanced beginner in about one day, using affordable off-the-shelf components (Table 1), and openly accessible data acquisition scripts running in Python and Arduino (Supplemental Files).

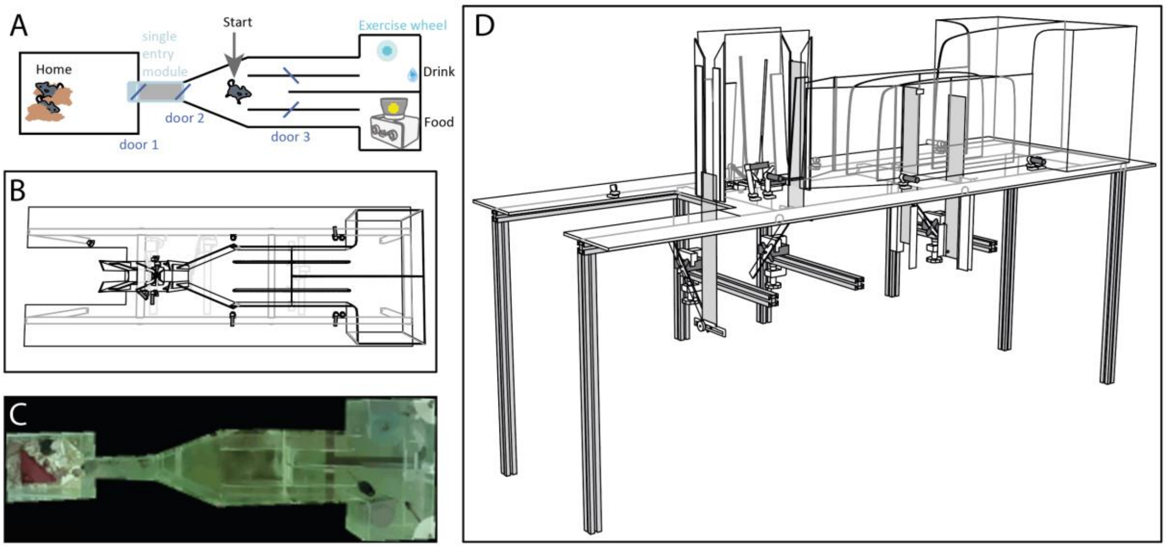


*Figure 1, Switchmaze. A, Schematic; B, CAD model from above; C, picture from above; D, CAD model.*

**Materials**

*Table 1, Bill of materials for the Switchmaze. Main parts for the three modules with separate build instructions in references[1–3] are labelled SEM (single entry module), RW (running wheel) and SWD (sensing water dispenser). 3d object file links are provided to enable simple modifications on www.tinkercad.com, while also .stl files are provided in supplementary files to allow simple printing.*

| Part | Count, vol. or length | Manufacturer/link | Manufacturer serial # or *.stl file | Approximate cost, EUR | Part | Module |
|---|---|---|---|---|---|---|
| Arduino Mega | 1 | Arduino | A000067 | 33.6 | | |
| Raspberry Pi 4b or 400 | 1 | Raspberry Pi | KW-2646 | 45 – 95.0 | | |
| Servo motor | 4 | Master | 1556176 - 62 | 112.0 | | SEM 2x, RW 1x |
| Small mirror tiles (1 x 1 cm), self-adhesive | 2 | e.g., amazon.com, or a piece of compact disc | N/A | 10.0 | | |
| Switch | 1 | Adafruit | P3064 | 2.9 | | |
| 5V power supply | 2 | RS Pro | 124-2183 | 23.0 | | |

| | | | | | | |
|---|---|---|---|---|---|---|
| wire | 2 m | RS Pro | 196-4225 | 0.8 | | |
| 2-core shielded cable | 20 m | Lapp | 0034302 | 10.0 | | |
| 3-core shielded cable | 2 m | Lapp | 1600207 | 6.0 | | SWD |
| 4-core shielded cable | 2 m | Lapp | 0034304 | 1.0 | | RW |
| RFID tags | 4 | Sparkfun | SEN-09416 | 22.0 | | |
| RFID tag implantation syringe | 1 | DHgate | 533480816 | 2.2 | | |
| 10 kOhm resistor | 6 | TE Connectivity | LR1F10K | 0.6 | | SEM 2x |
| Load cell 100 g | 1 | SparkFun | TAL221 SEN-14727 | 9.4 | | SEM |
| Open scale | 1 | SparkFun | SEN-13261 | 32.5 | | SEM |
| RFID antenna | 1 | SparkFun | ID-20LA SEN-11828 | 33.7 | | SEM |
| RFID USB reader | 1 | SparkFun | SEN-09963 | 26.3 | | SEM |
| Mini USB cable | 2 | SparkFun | CAB-13243 | 4.0 | | SEM |
| 650nm Laser Diode | 5 | SparkFun | P1054 | 28.0 | | SEM 2x |
| Photoresistor | 5 | SparkFun | SEN-09088 | 4.0 | | SEM 2x |
| Rotary encoder | 1 | Nidec Copal Electronics | REC16B50-201 | 20.7 | | RW |
| Transparent film A4 pieces | 2 | Staedtler | 636 10DT6F | 0.4 | | SEM |
| ‘flying saucer’ running wheel for small rodents | 1 | Ware pet products | #03281 | 7.0 | | RW |
| Rubber tubing 5mm ID/ 7mm OD | 20 mm | RS Pro | 235-4806 | 1.0 | | RW |
| Female to male jumper wire connectors | 25 | MikroElectronika | MIKROE-511 | 8.4 | | |
| FED3 | 1 | https://github.com/KravitzLabDevices/FED3 (or bought prebuilt from Labmaker or OEPS) | | 200.0 | | |
| Bio-Serv™ Dustless Precision Pellets™ for Rodents | 1 | Bio-Serv™ | F05684 | 162.0 | | |
| Relay module | 1 | TRU Components | TC-9927156 | 5.0 | | SWD |

| | | | | | | |
|---|---|---|---|---|---|---|
| Varistor 25VAC | 1 | TDK | B72205S0250K101 | 0.26 | | SWD |
| Resistor 10MOhm | 1 | Vishay | 594-CBB0207001003GCT | 0.6 | | SWD |
| Power source 24V | 1 | RS-PRO | 175-3306 | 10.0 | | SWD |
| Solenoid valve 24V | 1 | SMC | VDW12GA | 17.2 | | SWD |
| 50 ml syringe | 1 | Terumo | SS 50L1 | 5.4 | | SWD |
| 2.5 ml syringe | 1 | Terumo | SS 02LE1 | 0.5 | | SWD |
| 21 G hypodermic needle | 2 | Terumo | AN*2150R1 | 0.1 | | SWD |
| 3 mm (ID)/ 5mm (OD) silicone tubing | 100 mm | Saint Gobain | 760210 | 1.5 | | SWD |
| M5 threaded nylon barbed tube fitting | 2 | McMaster-Carr | 5463K557 | 1.2 | | SWD |
| Alligator clip | 1 | RS Pro | 738-5856 | 2.8 | | SWD |
| Chloroform* | 10 ml | Sigma-Aldrich | 288306 | 5.0 | | |
| 350 mm X 50 mm rectangles of 0.5 mm thick aluminium sheet | 4 | Reely | 297895 - 8J | 16.0 | door | SEM 2x |
| 3D-printed parts | 3 | https://www.tinkercad.com/things/0J1Yf0e4cs3-ball-joint-for-laser-httpswwwadafruitcomproduct1054 | bb345_send.stl https://osf.io/qrzsc | 20.0 all 3D-printed parts | A | |
| | 3 | https://www.tinkercad.com/things/h3bjfI7gLWr-bb1receive | bb345_receive.stl https://osf.io/dsjyr | | B | |
| | 1 | https://www.tinkercad.com/things/7VH506uKMcl-ladder | nest_ladder.stl https://osf.io/tbdq9 | | C | |
| | 2 | https://www.tinkercad.com/things/4CAZInAZsyC-doorstopper | HSD_door_stopper.stl https://osf.io/vtx2d | | D | |
| | 1 | https://www.tinkercad.com/things/aX0G8RGli2f-rwbaseandcover | rw_base.stl https://osf.io/p4hvf | | rwA | RW |

| | | | | | | |
|---|---|---|---|---|---|---|
| | 1 | https://www.tinkercad.com/things/aX0G8RGli2f-rwbaseandcover | rw_cover.stl https://osf.io/dx8mt | | rwB | RW |
| | 2 | https://www.tinkercad.com/things/gNGUW5mJAnq-copy-of-doorguidetop | HSD_door_guide_top.stl https://osf.io/vg693 | | E | |
| | 1 | https://www.tinkercad.com/things/4O52qQxFPTd-doorguidebottom | HSD_door_guide_bottomR.stl https://osf.io/w7uh3 | | F | |
| | 1 | https://www.tinkercad.com/things/4O52qQxFPTd-doorguidebottom | HSD_door_guide_bottomL.stl https://osf.io/dzpaq | | G | |
| | 1 | https://www.tinkercad.com/things/aOwZxV0AVPk-scale1 | scale1.stl https://osf.io/vq6xm | | semA | SEM |
| | 1 | https://www.tinkercad.com/things/3KBNiBqkAat-sembase | scale2.stl https://osf.io/sef5p | | semB | SEM |
| | 1 | https://www.tinkercad.com/things/3y3PaHMwj5M-semarches | scale3.stl https://osf.io/buv63 | | semC | SEM |
| | 2 | https://www.tinkercad.com/things/3y3PaHMwj5M-semarches | scale4.stl https://osf.io/4smzy | | semD | SEM |
| | 10 | https://www.tinkercad.com/things/8tfXQNYPWRZ-bbbase | bb_base.stl https://osf.io/c8fk9 | | H | SEM 4x |
| | 10 | https://www.tinkercad.com/things/8tfXQNYPWRZ-bbbase | bb_base_nut.stl https://osf.io/nmt4j | | I | SEM 4x |
| | 1 | https://www.tinkercad.com/things/1JqjB9udT5q-bb1send | bb1_send.stl https://osf.io/3tzwr | | semG | SEM |
| | 1 | https://www.tinkercad.com/things/h3bjfI7gLWr-bb1receive | bb1_receive.stl https://osf.io/he48d | | semH | SEM |

| | | | | | | |
|---|---|---|---|---|---|---|
| | 1 | https://www.tinkercad.com/things/2RNHwEzdBBe-bb2send | bb2_send.stl https://osf.io/r7hw2 | | semI | SEM |
| | 1 | https://www.tinkercad.com/things/gspdPT8vtrj-bb2receive | bb2_receive.stl https://osf.io/7c3t4 | | semJ | SEM |
| | 2 | https://www.tinkercad.com/things/co6F7bdX3kP-doorbase | door_base.stl https://osf.io/uef4y | | semK | SEM |
| | 2 | https://www.tinkercad.com/things/co6F7bdX3kP-doorbase | door_base_nut.stl https://osf.io/nmt4j | | semL | SEM |
| | 3 | https://www.tinkercad.com/things/5UCvPl0qUPy-servoclamp | servo_clamp.stl https://osf.io/n9mx5 | | J | SEM 2x |
| | 3 | | servo_bolt.stl https://osf.io/yvtjk | | K | SEM 2x |
| | 3 | | servo_bolt_cap.stl https://osf.io/kmh9s | | L | SEM 2x |
| | 1 | https://www.tinkercad.com/things/aPTybEE7aS5-rfidbase | rfid_base.stl https://osf.io/ngsfc | | semM | SEM |
| 3 mm Acrylic sheet parts | 1 | | middle_wall.svg | 50.0 all acrylic parts | M | |
| | 1 | | FED3_diagonal_wall.svg | | N | |
| | 1 | | FED3_base.svg | | O | |
| | 2 | | FED3_structural_support.svg | | P | |
| | 1 | | floor.svg | | Q | |
| | 1 | | HSD_door_lever.svg | | R | |
| | 4 | | inner_wall.svg | | S | |
| | 2 | | long_side_wall.svg | | T | |
| | 2 | | connecting_side_wall.svg | | U | |
| | 2 | | goal_wall1.svg | | V | |

| | | | | | | |
|---|---|---|---|---|---|---|
| | 1 | | nest_back_wall.svg | | W | |
| | 1 | | nest_back_wall_water_base.svg | | X | |
| | 2 | | nest_back_wall_water_base_support.svg | | Y | |
| | 1 | | nest_front_wall_connector.svg | | Z | |
| | 2 | | nest_front_wall_half.svg | | AA | |
| | 2 | | nest_long_wall.svg | | BB | |
| | 2 | | goal_wall2_side.svg | | CC | |
| | 1 | | goal_wall3_back.svg | | DD | |
| | 4 | | door_guide_back.svg | | semN | SEM |
| | 4 | | door_guide_spacer.svg | | semO | SEM |
| | 4 | | sem_support.svg | | semP | SEM |
| | 2 | | door_lever.svg | | semQ | SEM |
| 1 mm Acrylic sheet parts | 4 | | door_guide_front.svg | | semR | SEM |
| | 2 | | prevent_burrowing.svg | | semS | SEM |
| 20 mm square profile aluminium rail, 500 mm length | 4 | McMaster-Carr | 5537T101 | 132.2 all rail parts | | |
| 20 mm square profile aluminium rail, 300 mm length | 9 | McMaster-Carr | | | | SEM 4x |
| 20 mm square profile aluminium rail, 1400 mm length | 2 | McMaster-Carr | | | | |
| M6 cap screws, 30 mm length | 6 | McMaster-Carr | 90128A266 | 9.2 | | |
| M6 grub/set screws, 30 mm length | 10 | McMaster-Carr | 92015A136 | 6.5 | | SEM 4x |
| M6 Drop-In T-Nut | 10 | Thorlabs | XE25T1/M | 30.5 | | SEM 4x |
| M3 cap screws, 30 mm length | 4 | McMaster-Carr | 91290A171 | 7.2 | | SEM 2x |

| | | | | | | |
|---|---|---|---|---|---|---|
| M3 cap screws, 10 mm length | 5 | McMaster-Carr | 91274A105 | 6.4 | | SEM 2x |
| M3 nuts | 6 | McMaster-Carr | 90592A085 | 2.6 | | |

Approx. total cost: 1137 – 1187 EUR

* Welding acrylic and 3D-printed PLA parts with chloroform must be done safely in a well-ventilated space, with minimal amounts of chloroform and using appropriate PPE including gloves and an appropriate mask.

The 3D-printed and laser cut part drawings can be found at https://osf.io/yb67q/ and https://github.com/MaheshKarnani/Switch_maze/tree/main/Modules_SM/Switchmaze.

Useful tools: drill, Allen key set, a small screw driver, wire cutters, soldering iron, tape, 1 ml syringe for the chloroform, personal protective equipment and epoxy glue.

#### Build instructions

##### General description

The Switchmaze incorporates six modules: the single entry module (SEM)[1], sensing water dispenser (SWD)[3], timed running wheel[2], a FED3 pellet dispenser[4], a horizontally sliding door (HSD) and a home cage with added walls. These parts are embedded in a simple maze made of 3 mm thick acrylic sheet built on an elevated 1400 by 500 mm floor plate (Figure 1D, interactive 3D model in Supporting Files).

Most parts are attached to each other using chemical welding with chloroform. This is a hazardous technique which requires adequate ventilation and PPE including gloves and an appropriate mask. Using minute quantities of chloroform minimizes risks, so it is strongly recommended to dispense chloroform sparingly from a 1 ml syringe (blunted needle for safety). The general idea is to firmly hold two pieces of acrylic sheet (or an acrylic sheet and 3D-printed PLA component) against each other and allow a drop of chloroform to seep in between the pieces and keep holding them in place for about one minute. After this, the bond will have formed and most of the free chloroform will have evaporated. If needed, the bond can usually be broken by bending the pieces, and remade, as long as the pieces have enough contact surface with each other. Acrylic cement (e.g., Acrifix 192) may be used to form permanent, cavity-free bonds after the initial weld.

##### Mechanical assembly

1. ***Frame and floor plate:***

A support frame for the floor plate is built using three 300 mm, two 1400 mm and four 500 mm long 20 mm square profile aluminium rail according to Figure 2 and lay the 3mm acrylic sheet floor plate on top (part Q). A 6 mm drill piece suitable for aluminium is used to make holes through the 1000 mm rails for the 300 mm cross beams. Steel 6 mm screws will readily form a thread in the axial hole of the rails when first screwed in, but this is best done before assembly as some force may be required. Drop-in nuts and grub screws are used for the 500 mm legs. A minimum height of 500 mm from the ground is recommended as the doors need to operate some distance below the floor plate.

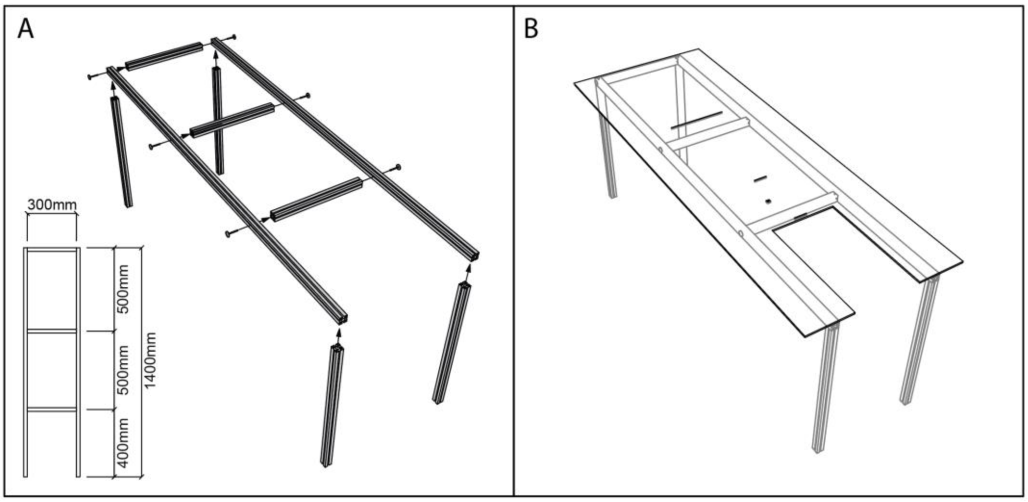


*Figure 2, Aluminium rail frame without (A) and with floor plate (B).*

**2. *Single entry module:***

The SEM is assembled on the floor plate (Figure 3), using parts labelled SEM in the 'Module' column of Table 1. It consists of a 200 mm long U-shaped corridor, constructed from a sheet of transparent, firm, light-weight plastic (made from two overhead projector transparency sheets) which is secured into a U-shape via a 3D-printed skeleton (PLA). The U-shaped corridor is screwed onto a 100 g load cell (SparkFun TAL221 SEN-14727) which is secured to the floor plate. The load cell is connected to a USB amplifier (SparkFun SEN-13261). Laser cut 3 mm acrylic sheet pieces are used to construct edge guides for the two doors by chemical welding using chloroform (using appropriate PPE). Additional pieces of acrylic sheet are attached to the door edge guides to support the edges of the U-shaped corridor in case an animal pushes against it. The SEM can be closed off on both ends via vertical sliding doors constructed from 300 mm X 50 mm rectangles of 0.5 mm thick aluminium sheet which can be lowered into the floor of the setup via a servo motor attached to a beam below. An RFID-tag reader (Sparkfun ID-20LA and SEN-09963) is placed below the U-shaped corridor, near door 2. Two beam break devices (BB), constructed from a 650 nm laser diode (SparkFun P1054) and photoresistor (Sparkfun SEN-09088) circuit mounted on 3D-printed ball joints are used to ensure safe operation of the doors. One (BB1) is directed 10 mm above door 1 in open position. It is used to prevent door 1 closing with an animal on top. This beam is unbroken when door 1 is open and no animal is sitting in the opening. Another (BB2) is directed through the corridor at a safe distance (>100 mm) from door 2. A break in this beam signals exit through the module back to the home cage.

Before beginning, parts semA-semM and H-L should be printed (Figure 4).

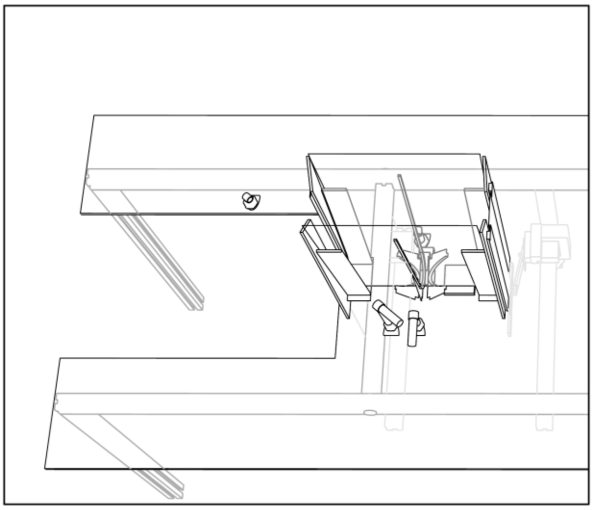

*Figure 3, Single entry module.*

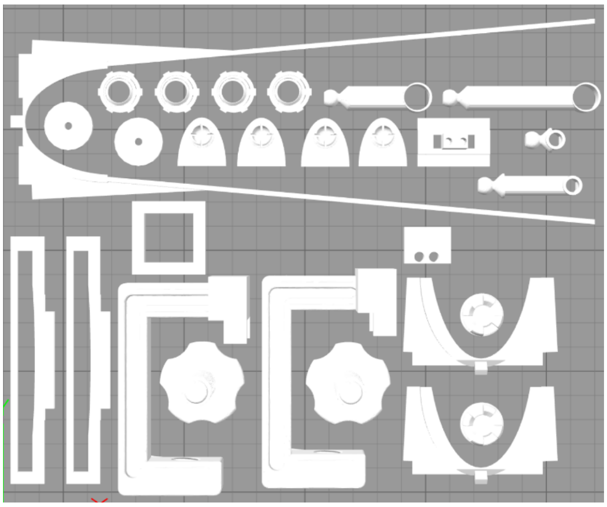

*Figure 4, 3D printed parts for the SEM.*

A) U-shaped corridor with RFID detector and scale:

The door guide frames of the U-shaped corridor are assembled using chemical welding of acrylic sheet pieces. This is a hazardous technique which requires adequate ventilation and PPE including gloves and an appropriate mask. Using minute quantities of chloroform minimizes risks, so it is strongly recommended to dispense chloroform sparingly from a 1 ml syringe with a blunted needle for safety.

Four guide frames for the doors are constructed using acrylic sheet pieces semN, semO and semR (Figure 5A). Each set of three pieces is clamped together with a binder clip (Figure 5B) and a drop of chloroform is used to bond them, making sure the three pieces are aligned at the bottom for a strong bond to the floor plate. Each frame forms a 3 mm slot that functions as a rail for one side of a door. Care should be taken to align these slots with the 3 mm door aperture in the floor plate, and to leave enough room for the door to slide up and down while remaining within the guide frame (Figure 5C). The process is repeated on the other edge of the door, and twice more for the frames of the second door. It is useful to confirm the fit with a ‘door’ aluminium sheet piece, which should be loosely within the frame (Figure 5B,C).

The RFID detector (SEN-11828 connected to SEN-09963) is attached, with the antenna (ID-20LA, SEN-11828) facing upward, to the square shaped 3D-printed support (semM) using two small drops of superglue, and this assembly is placed near door 2, leaving a minimum of 25 mm clearance between the door frame and RFID detector (Figure 5D). The support can then be joined to the floor plate with a drop of chloroform.

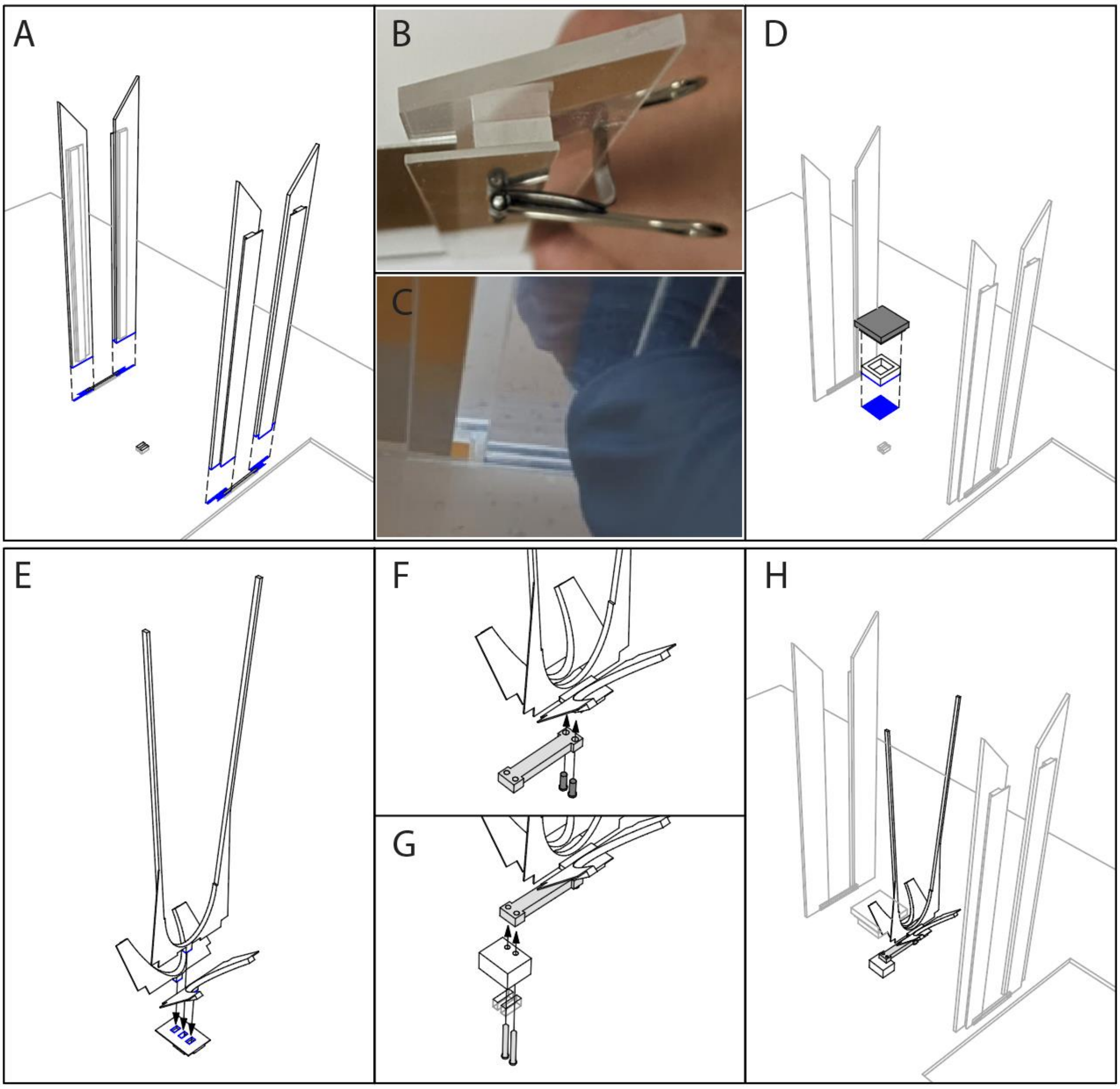


*Figure 5. Guide frame, RFID detector and scale backbone installation. Use PPE and minimal amounts of chloroform.*

After guide frames for both doors are built, the U-shaped corridor with the scale is assembled as follows. A scale backbone is assembled from 3D-printed scale support (semA-semD) and a 100 g load cell (Sparkfun SEN-14727) by push-fitting the pieces as shown in Figure 5E and securing it to the loadcell using M3 screws (Figure 5F). The prints may need to be filed down at the attachment points to get a good fit. A loose push-fit is acceptable at this stage as a solid joint will be made with epoxy-glue at a later stage. The complete scale backbone is screwed in place firmly with two 30 mm M3 screws passed through the 10 x 3 mm slits in the floor plate and 3D-printed part semA. The screws are tightened into the threaded holes on the load cell (Figure 5G).

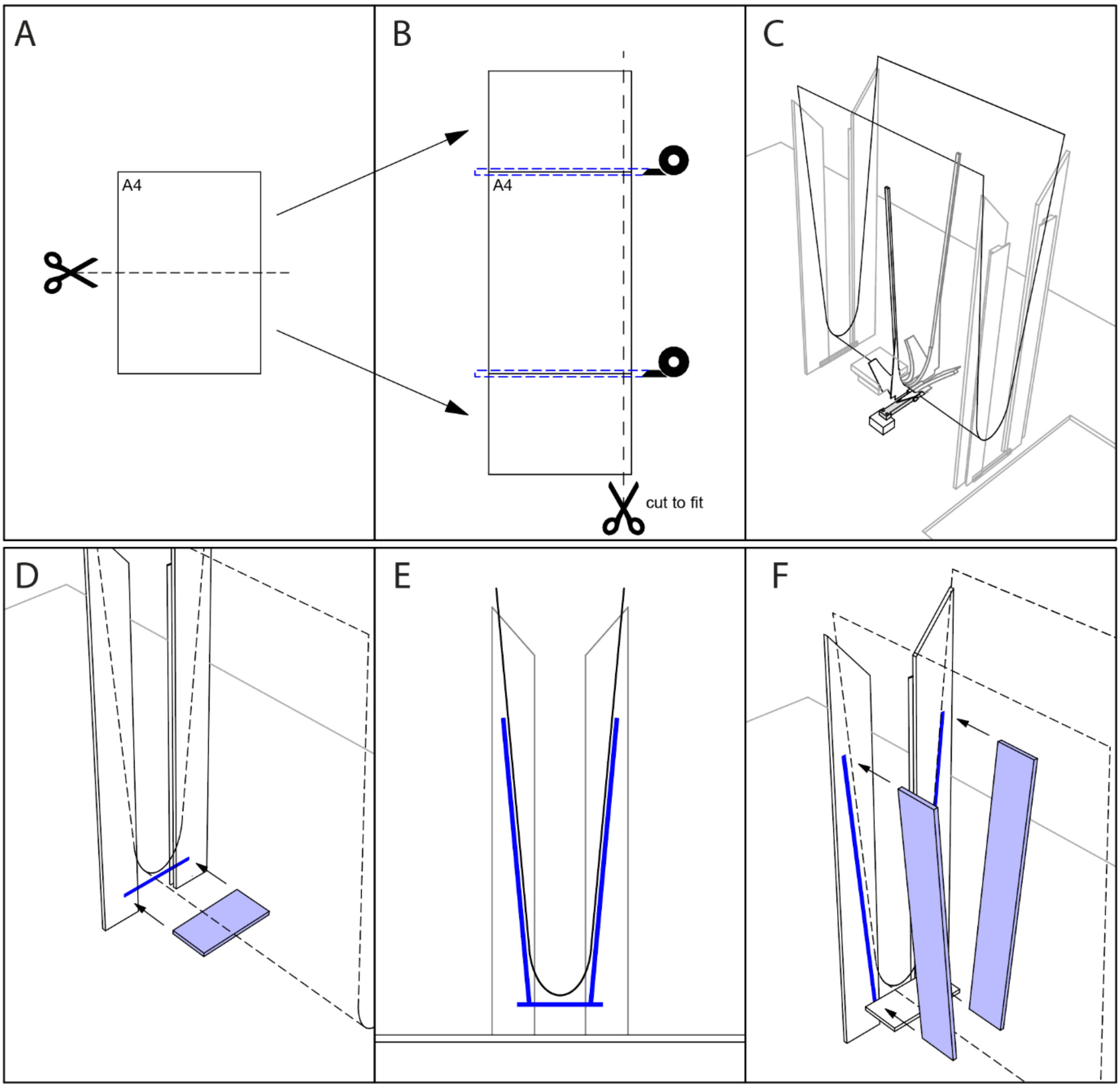


*Figure 6. U-shaped corridor and support structures.*

A 597 x 200 mm rectangle of transparency film is made from two A4 sheets of film, cutting one in half and taping the halves to the ends of the intact one, followed by cutting 10 mm off the long edge (Figure 6A,B). The film is then bent into a U-shape that fits inside the scale backbone and taped in place along the tall middle supports. It may be necessary to cut the ends slightly to make the film fit snugly in between the doors (Figure 6C). The edges of the film should not push against the door frames, but there should also be no gaps larger than 1-2 mm. A new piece of film is simple to install at this stage if the gaps are too large after cutting. When an acceptable fit is achieved, the film is taped firmly in place along the tall middle supports of the backbone and epoxy glued to the backbone along the U-shaped parts hugging it. 1-2 ml of epoxy should be applied to firm up any loose push-fit junctions between the 3D-printed parts. The finished U-shaped corridor should be a solid but light-weight piece that transmits the animal's weight to the load cell regardless of animal movement in the scale.

Next, two 1 mm thick acrylic pieces (semS), that prevent burrowing below the scale, are attached using a drop of chloroform to the door frame guides about 2 mm below the transparent film (Figure 6D).

Then, four 3 mm thick acrylic pieces (semP) are attached to the door frame guides using drops of chloroform, about 1 mm away from the transparent film (Figure 6E,F). The film may lean against these supports toward the top, but not at the bottom. This way the animal's weight will be measured accurately while the film wall appears sturdy enough to discourage excavation.

B) Doors:

To prepare the moving parts (Figure 7), a lever made of 3 mm acrylic sheet (semQ) is tightly attached to a servo hub (provided with the servo) using a 10 mm M3 screw. In a well-ventilated space/chemical hood, 1-2 drops of chloroform are added onto the joint to activate the surfaces. After a few minutes 2-3 drops of superglue are added onto the joint. After the glue has set, the M3 screw can be removed. A 300 mm X 50 mm rectangle of 0.5 mm thick aluminium sheet (part "door") is slotted into a 3D printed base (semK) and 2-3 drops of superglue are added onto the joint. A small amount of epoxy glue can be used to bond these glued joints as they may need to carry high loads if installed at less than ideal angles or if the unit is not kept sufficiently clean. A 30 mm M3 cap screw is secured firmly with a M3 nut to the free end of the lever (see also Figure 8C).

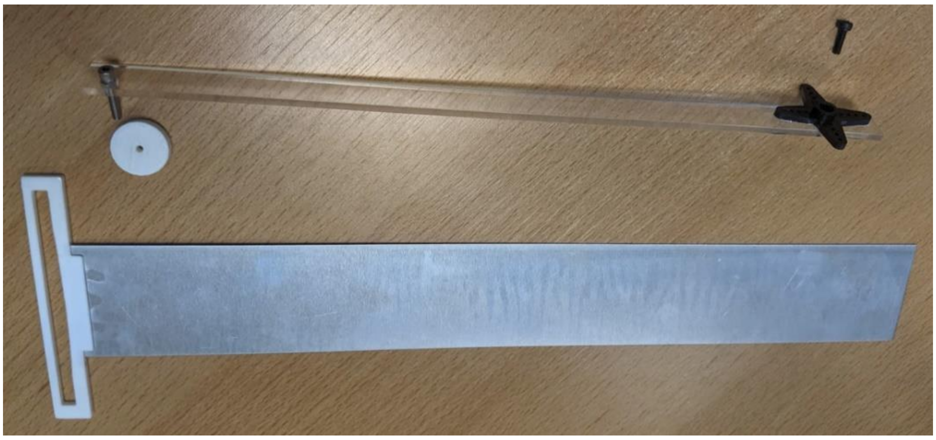

*Figure 7. acrylic door lever along with attachment pieces (above) and aluminium door (below).*

Using the drop-in T-nuts and M6 grub screws, a 300 mm aluminium rail is hung downward from the support frame, and another 300 mm aluminium rail is attached perpendicular to that (Figure 8A). A servo motor is positioned on top of the perpendicular rail and secured using 3D printed clamp (parts J-L; Figure 8B). The door lever to the servo hub is attached such that its movement arc cannot collide with the floor, i.e., so the 180 degree range of the servo stops before the lever would collide with the floor. The door is inserted through its aperture in the floor and the 30 mm screw of the lever is threaded through the door's base (Figure 8C). Then the screw is capped using the 3D printed nut (part L), but not tightened against the door base (see Figure 8C inset). While the servo is powered off, the door can be tested by manually moving the lever up and down (Figure 8D-F). The lever should operate perpendicular to the door, such that the only part touching the door is the 30 mm screw. If the lever's distance to the door base changes during operation, the servo may need to be aligned better by opening the 3D printed clamp, readjusting the servo position and closing the clamp again. There should be minimal friction during door operation and the door should remain in the frame slots during the up and down motion. If strong friction is noticed, the components can be removed (welded joints can be snapped off) and reinstalled at slightly altered locations. Lubricant use should be minimized as the animals can spread it into the rest of the apparatus, however, minimal amounts of Vaseline can be applied to the 30 mm screw shank and the edges of the door.

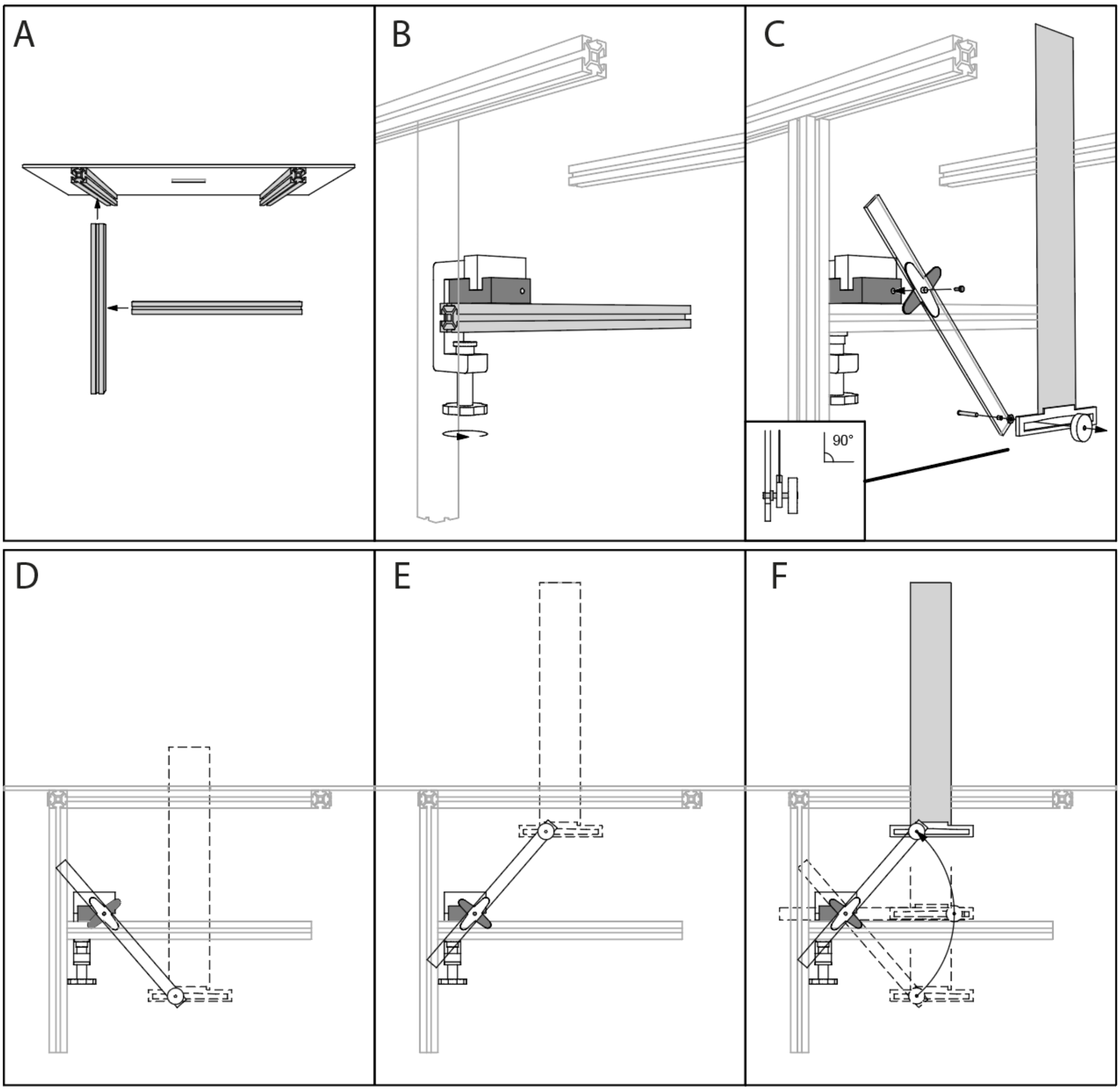


*Figure 8. Door installation.*

C) Beam break detectors:

Lastly, the beam break detectors are installed using 3D-printed holders (parts H-I and semG-semJ), diode lasers (Sparkfun P1054) and photoresistors wired to resistors. The sender and receiver holders are built first (each has 3 parts: base H, tightening nut I and one of the parts semG-semJ with a ball joint; Figure 9). The ball joint may need to be smoothed using sandpaper, depending on the grain of the print.

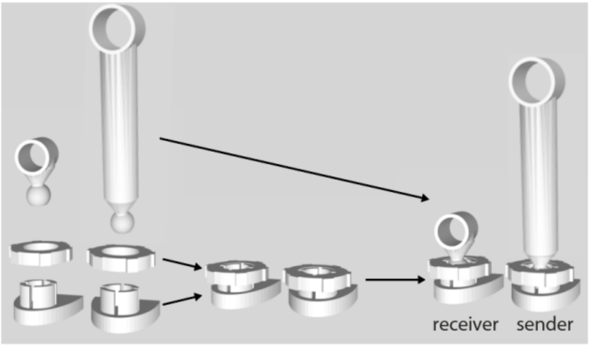

*Figure 9. Beam break detector holder assembly.*

After assembly, the parts are placed at their approximate locations (Figure 10) and door 1 is set to protrude approximately 50 mm above the floor plate (closed position). Then beam targeting is checked by sighting through the laser holders to the detectors. Beam 1 needs to pass in the middle of the entrance about 10 mm above door 1 (Figure 10A,C). Beam 2 needs to pass about 10 mm above the bottom of the U-shaped corridor at a distance of >100 mm from door 2 (Figure 10A,B). A small drop of chloroform is used to weakly bond each holder to the floor plate, so they can be detached if necessary. Later when the lasers and doors are operational, locations can be confirmed and the holders' base parts should be joined strongly to the floor with additional drops of chloroform.

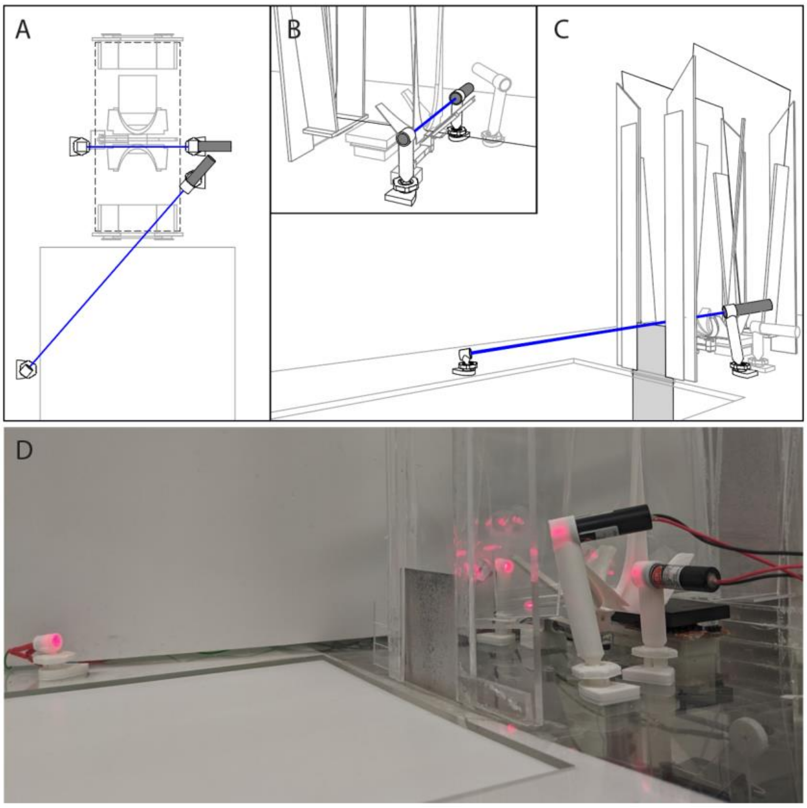

*Figure 10. Beam break (BB) devices. BB1 aimed across the top of door 1, and BB2 aimed perpendicularly through the U-shaped corridor.*

At this stage the SEM mechanical build is complete and should look like Figure 3.

3. ***Walls of the foraging environment:***

Next, building the maze should be continued from the SEM toward the reward zone. All walls are installed by chemical welding using a few drops of chloroform and gently pushing the bottom edge of the wall against the floor plate for about one minute while the bond forms. Two heavy objects with a straight edge can useful, as the wall can be sandwiched in between them and allowed to bond with the floor for some minutes. It is best to use a ruler and a marker pen to draw in the wall positions first. After joining to the floor, each wall should be bonded to an adjacent wall, except for the four inner walls (see below). Walls should be joined in the following order:

A) First, the long side walls (part T, 400 x 200 mm) whose middles (400 mm long side) should align with the ends of the 2 x 200 mm HSD hole in the floor plate (Figure 11A).

B) Then the two connecting side walls (part U, 200 x 200 mm) should connect diagonally the long side walls and the SEM door frame (Figure 11B).

C) Then the five walls of the reward area, by sequentially attaching parts V (110 x 300 mm with one round corner), CC (200 x 300 mm) and DD (400 x 300 mm) to the ends of the long side walls (Figure 11C). The final positions of these pieces may need to be adjusted so it is best to use small amounts of chloroform and weld them initially in quick succession so they can be moved if necessary. When final positions are reached, a strong bond should be formed between adjacent walls and the floor using a few drops of chloroform and firm force on each joint.

D) The middle wall (part M, 400 x 300 mm with one round corner) is added to separate the two goal areas (Figure 11D).

E) The four inner walls (part S, 200 x 200 mm with one round corner) are added at a roughly 50 mm distance from the side walls to separate entry and exit passages to/from the goal areas (Figure 11E,F). Care must be taken to keep the inner walls away from the 2 x 200 mm HSD aperture (Figure 11G) as the HSD will slide horizontally through the gap between these walls.

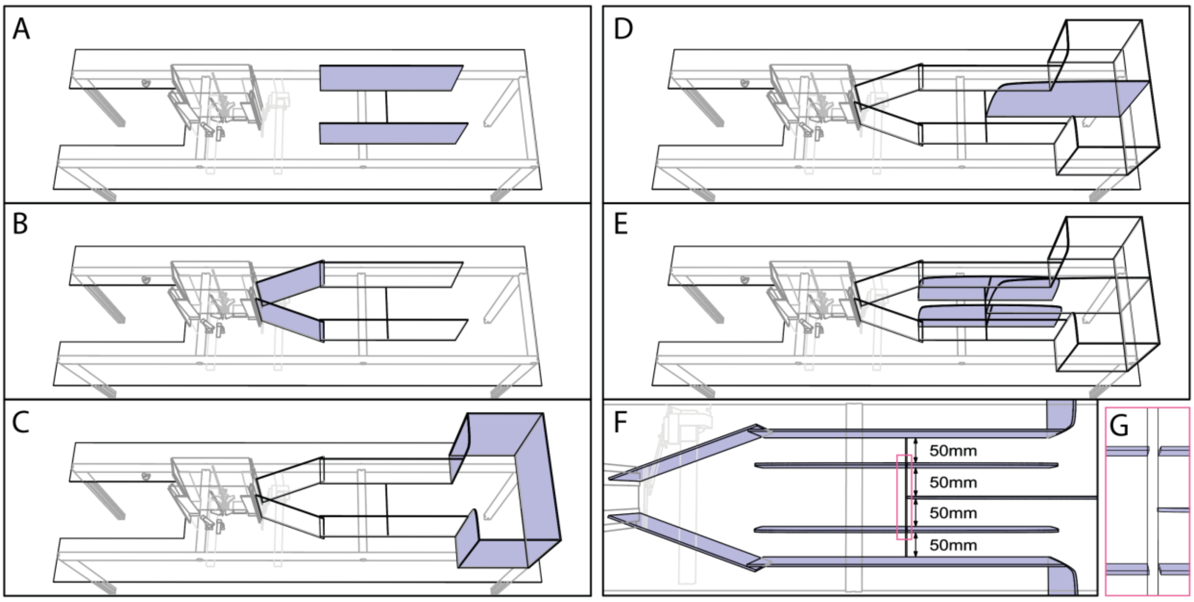


*Figure 11, Walls of the foraging environment.*

**4. *Horizontal Sliding Door:***

Using the drop-in T-nuts and M6 grub screws, a 300 mm aluminium rail is hung downward from the support frame, and another 300 mm aluminium rail is attached perpendicular to that (Figure 12A). A servo motor is attached on top of the perpendicular rail using a 3D-printed clamp (parts J-L; Figure 12B). The hub of the servo should be aligned with the middle wall (Figure 12D).

The HSD lever is glued to a servo hub attachment: The HSD lever made of 3 mm acrylic sheet (part R) is attached at its middle hole to a servo hub attachment with a flat surface (provided with the servo) using a 10 mm M3 screw. Once tightened, working in a well-ventilated space/chemical hood, 1-2 drops of chloroform are added on the joint to activate the surfaces. After a few minutes 2-3 drops of superglue are added to the joint. After the glue has set, the M3 screw can be removed. Epoxy glue is used to reinforce the joint as it will need to withstand impacts in routine operation. The adhesives should be allowed to set completely such that the lever and hub attachment are held together strongly.

3 mm holes are drilled through the aluminium doors (part 'door'; 50 x 350 mm aluminium sheet) at the indicated locations (Figure 12C). Then, the 3D-printed bottom side door guides (part F and G) are attached by sliding them on the edges of the doors as indicated and gluing in place with small drops of superglue (Figure 12C). These door guides will oppose each other when the door is in the closed mode (middle passage closed).

The doors can then be assembled by fastening the lever to the servo hub, making sure that the travel range is appropriate for door operation (the servo should be at its end of range when the lever is nearly touching the floor plate at near vertical angle), and attaching the doors with 10 mm M3 cap screws and nuts to the ends of the lever (Figure 12E). These screws should not be tightened all the way as the angle between the lever and doors changes during operation.

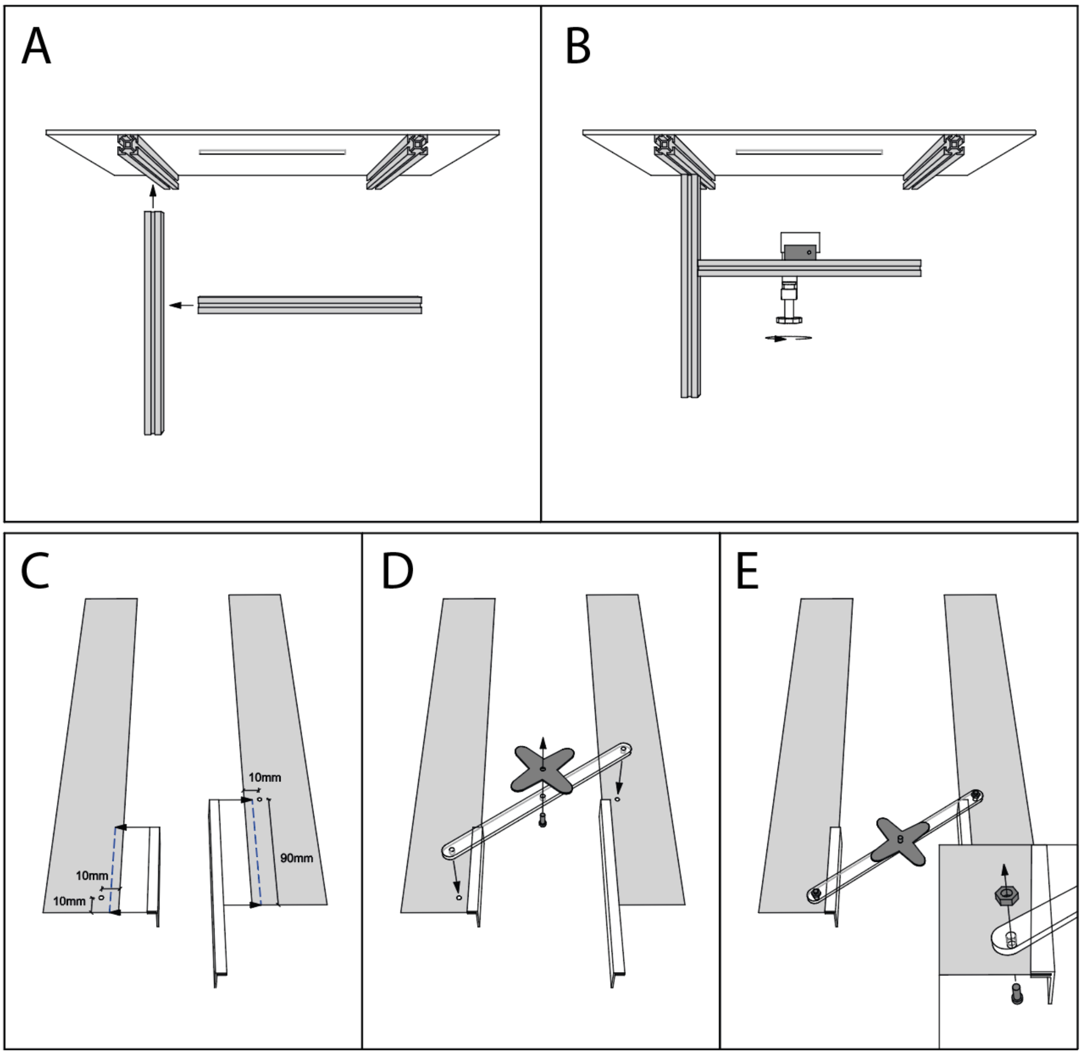


*Figure 12, Assembling components of the HSD.*

Finally, the limiting stoppers and top guide piece are attached to the doors (Figure 13A). The top door guide (part E) is glued to the top of one door (Figure 13B), such that it will oppose the middle wall when the door is closed. To fit the limiting door stoppers (part D), the doors must first be moved to the fully open position (Figure 13A). This can be done manually as long as the servo is not powered. The door stoppers should be welded to the floor plate with a small amount of chloroform such that they touch the doors when they are in the open position, forcing them to be vertical (Figure 13C,D). These stoppers function to keep the doors vertical when they are in the open position (middle passage open), and therefore should make contact with the aluminium doors in the open position (Figure 13D). The top and bottom door guides function to keep the doors vertical in the closed position (Figure 13E).

When the door is in operation, the angle commands should be selected carefully so that the stoppers and guides keep the doors vertical but no force is left acting on the doors at open or closed angle, as this will damage the components. I.e., the servo should have free range to move a couple of degrees beyond the open position. Similarly, in the closed position, the doors should not generate a static force against each other, but should just right each other.

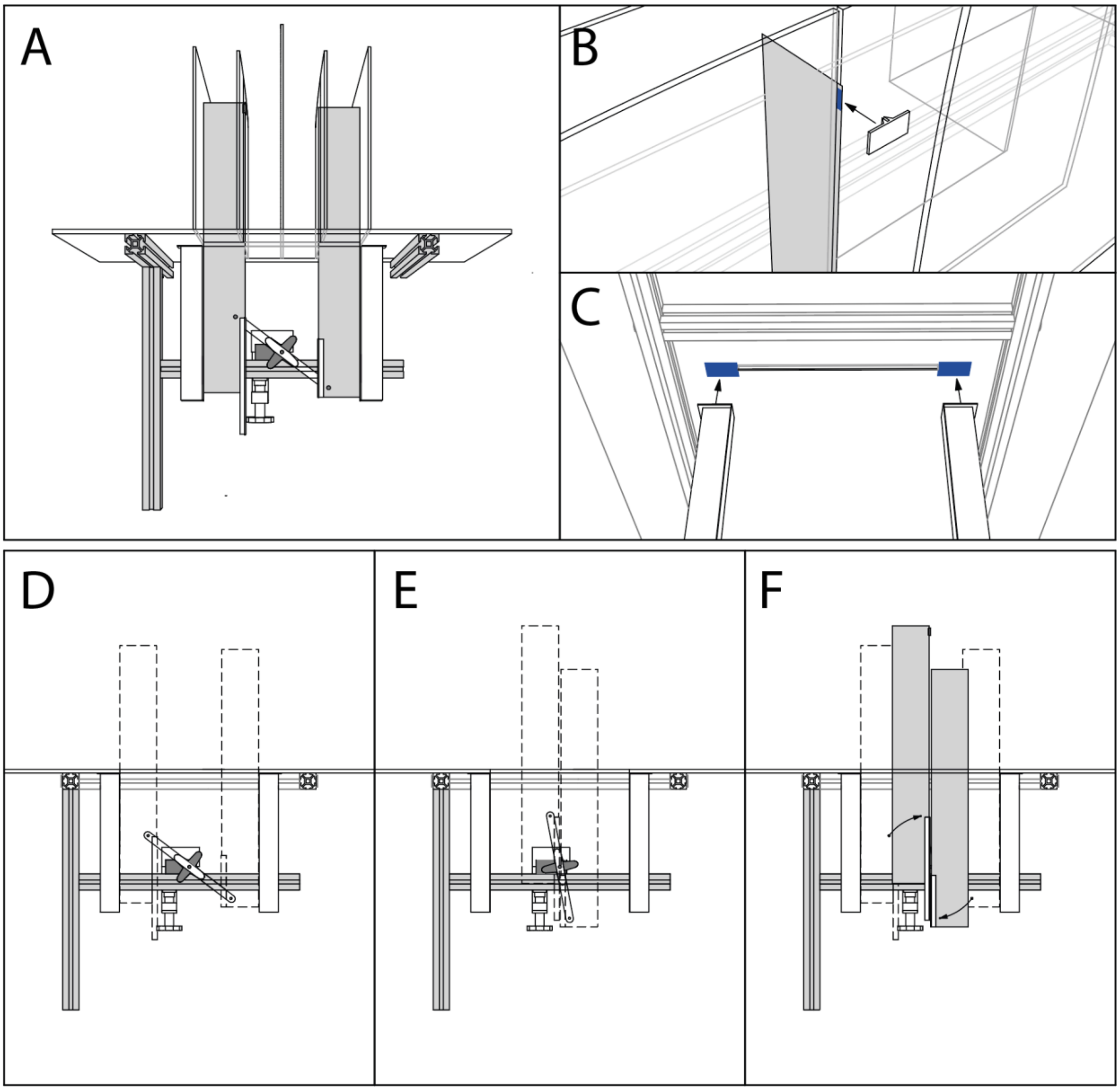


*Figure 13, Horizontal sliding door with stoppers and guides.*

**5. *Beam break detectors*:**

Three individual beam break detectors are built from identical components using 3D-printed parts A,B,H and I (Figure 14A), diode lasers (Sparkfun P1054) and photoresistors wired to resistors. The laser and detector holders are built first (Figure 14A). Each has three parts: base (part H), tightening nut (part I) and a part with a ball joint (part A for the laser and part B for receiver). The ball joint may need to be filed smooth depending on the grain of the print. After assembly, the parts are placed at their approximate locations (Figure 14B) and a thin mirror, such as a piece of a compact disc is glued or welded to the middle wall so beams from senders 3 and 4 will be reflected to their detectors. Beams should always be about 10-15 mm from the floor to ensure animal detection. Beam targeting is checked by sighting through the senders to the detectors. A small drop of chloroform is used to weakly bond each holder to the floor plate, so they can be detached if necessary. Then the lasers and photoresistors are inserted into their holders (use small abount of glue if needed). The cylindrical part of the receivers should be printed using black plastic or coated with a black paint (e.g., black nail polish) or black tape so that the major source of detected light is the sender. Later when the lasers are operational, locations can be confirmed and the holders' base parts should be joined strongly to the floor plate with additional drops of chloroform. The ball joints allow fine tuning the beam targeting.

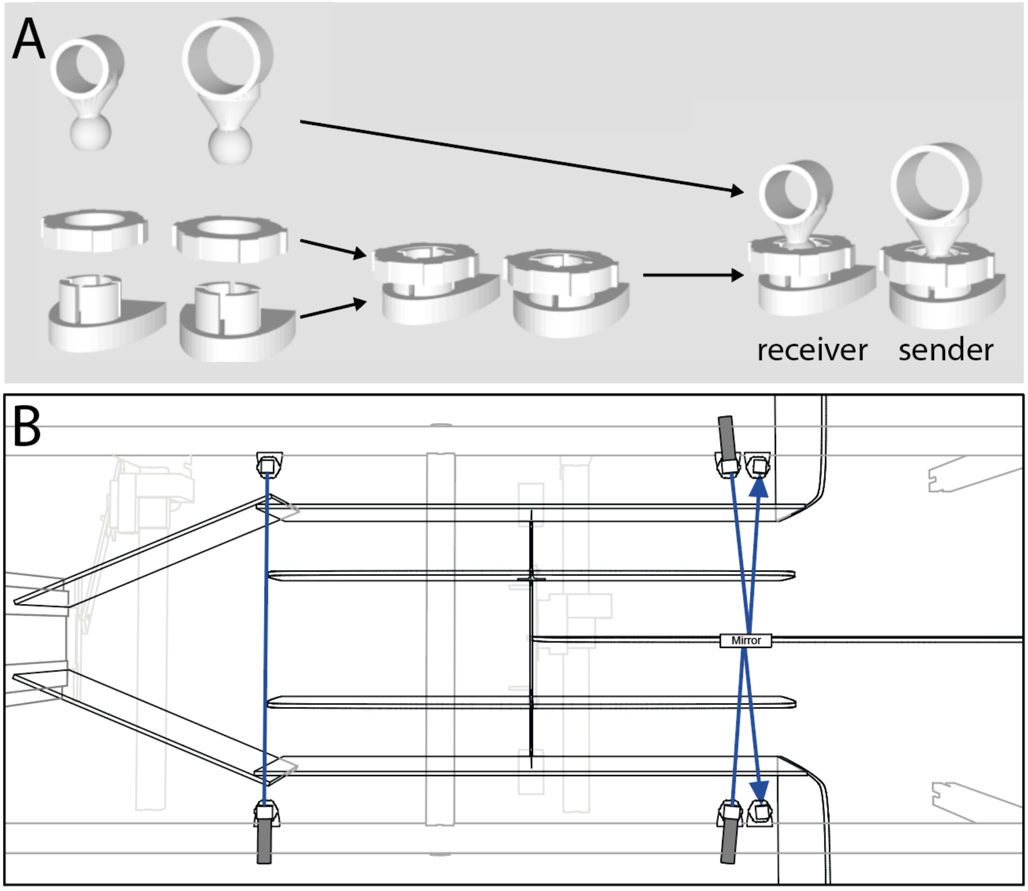


*Figure 14, Beam break detectors. A, Each sender and receiver unit is assembled from three 3D-printed parts. B, Beam break detectors are placed at a safe distance (>100 mm) from door 3. For detectors 4 and 5, a thin mirror, such as a piece of a compact disc, is glued to the middle wall.*

At this stage the device should look like Figure 1B and D.

#### 6. *Home-cage:*

The home-cage bottom of each cohort of mice is used as the home area in the Switchmaze (Figure 15A). It has a top edge protrusion 'lip' (Figure 15B) which allows it to slide into the 200 mm wide slot on the floor plate.

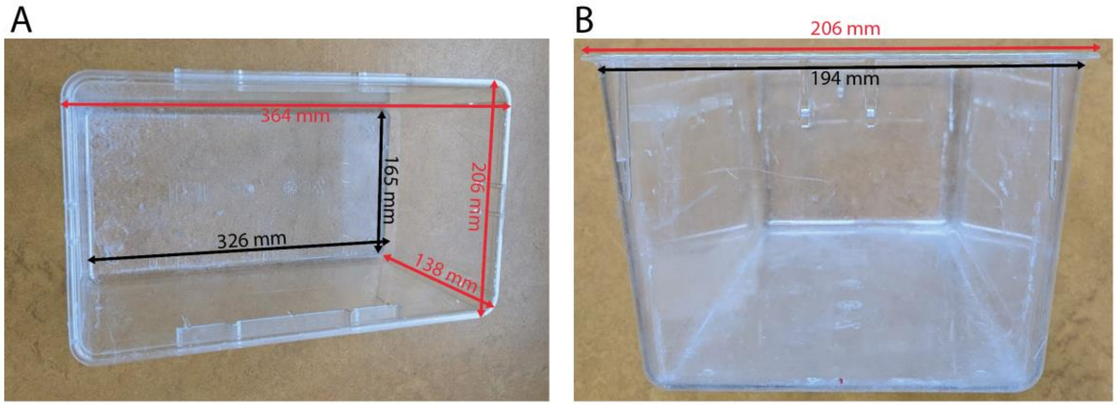


*Figure 15, Home-cage bottom.*

A wall unit for the home-cage is built from 3 mm thick acrylic sheet and a 3D-printed ladder (parts C, W, X, Y, Z, AA, BB). These are welded together with chloroform sequentially as shown in Figure 16.

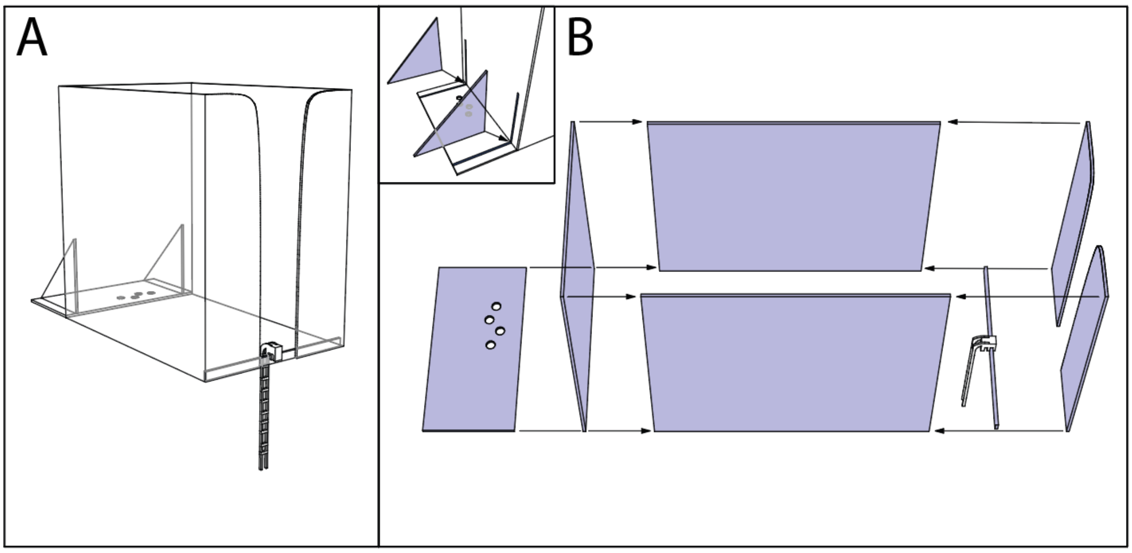


*Figure 16, Home wall assembly.*

#### 7. *Goal modules:*

Food, water and a running wheel are provided as goal modules that can be easily swapped between the goal areas, removed or replaced with other items, such as social chambers or novel objects.

A) Sensing water dispenser:

A sensing water dispenser (SWD, Figure 17) is assembled using parts labelled SWD in the 'Module' column of Table 1. The Arduino Mega performs capacitive sensing of the water spout to sense licking and transmits detected licks to the Raspberry Pi which saves data and controls water availability through a high-end switch (TRU Components TC-9927156) connected to a solenoid valve (SMC VDW12GA).

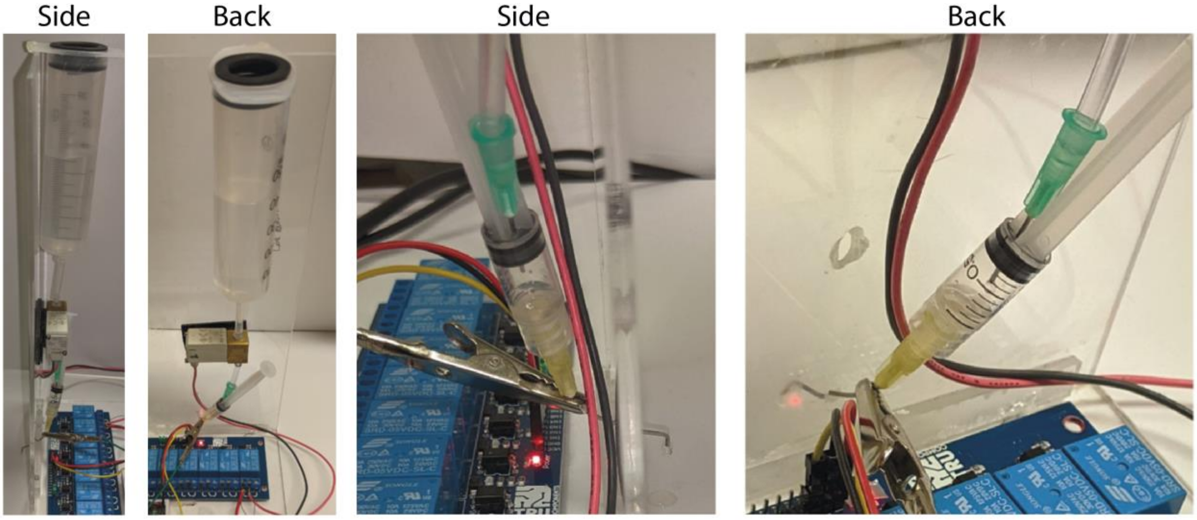


*Figure 17, Side and back views SWD, and magnified pictures of the isolated small reservoir sensed by capacitive measurement (two right-most panels).*

The water reservoir - a 50 ml plastic syringe - is connected, using 3 mm ID, 5mm OD silicone tubing and a M5 threaded nylon barbed tube fitting (McMaster-Carr 5463K557), to a solenoid valve. From the valve, 30 mm of the tubing is connected to a 21 G needle (forcing the 5mm OD tube inside the 4mm ID luer lock forms a strong junction) which is inserted through the plunger and rubber stopper of a shortened 2.5 ml syringe so that upon valve opening water will drop into an isolated small reservoir in the syringe (Figure 2). This smaller syringe acts to isolate the large reservoir of water from the water inside the lick spout which is sensed during capacitive measurement. The 2.5 ml syringe itself is fitted with a blunted, bent 21 G needle, which functions as the licking spout. An alligator clip connects to the licking spout to sense capacitance changes during licking. To ensure that there is no contact between the spout and the maze itself, the middle of the needle can be insulated with 20 mm of rubber tubing, leaving the tip of the spout exposed. The spout is bent downward so hanging drops do not spread along the spout changing its capacitance.

The water spout is introduced through a 4 mm diameter hole drilled in the wall at about 10 mm above the floor plate (Figure 18A). The rubber stopper of the 50 ml syringe can be used as a dust cap if a small hole is made in it to prevent pressure build up

B) FED3:

A Feeding Experimentation Device, FED3[4] is built following Kravitz lab instructions https://github.com/KravitzLabDevices/FED3 (or bought prebuilt) and modified for simultaneous input and output:

i. A 3-core cable is soldered on the FED3 main PCB front side such that it carries the BNC output signal lead, ground and Feather M0 Adalogger pin 9.
ii. The FED3.cpp code (December 2020 version) is modified by changing line 352 from `pinMode(BNC_OUT, INPUT_PULLDOWN);` to `pinMode(BNC_IN, INPUT_PULLDOWN);` and line 354 from `if (digitalRead(BNC_OUT) == HIGH){` to `if (digitalRead(BNC_IN) == HIGH){`.
iii. The FED3.h code (December 2020 version) is modified by adding `#define BNC_IN                    9` between lines 58 and 59.
iv. After these changes, the 'Dispenser' code is flashed to the FED3.

With these modifications, the FED3 will dispense a pellet when the Feather M0 Adalogger pin 9 goes high and will report a pellet retrieval on the usual output BNC port (A0).

The FED3 is placed in a diagonal wall assembly that is chemically welded together using parts N and O and two copies of part P (Figure 18B-D). This structure makes it easy to take out the FED3 for cleaning and protects its cables.

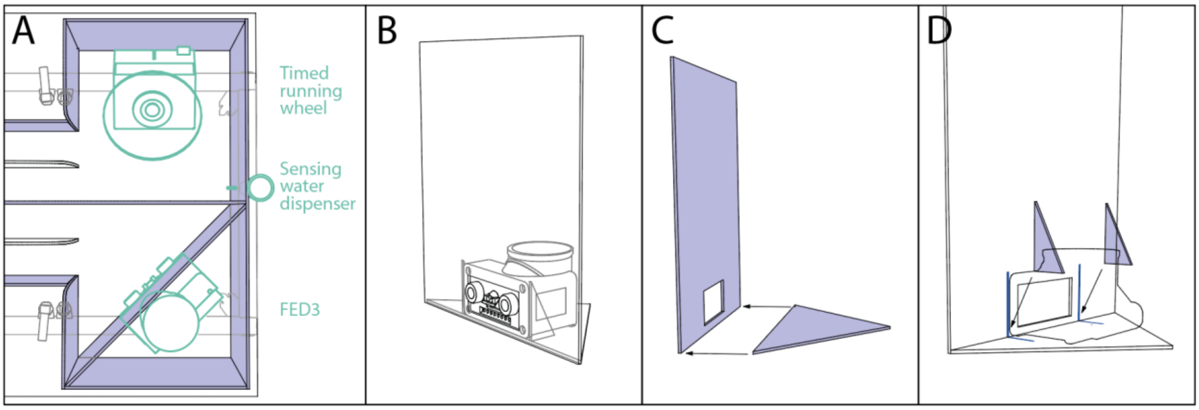


*Figure 18, goal modules. A, placement of goal modules (cyan). B-D, FED3 module assembly.*

C) Running Wheel:

A timed-access running wheel (RW, Figure 19) can be built using parts labelled RW in the 'Module' column of Table 1. However, given the low use of the running wheel in Switchmaze (see below and main document), it can be left out of the build. The Arduino Mega controls a servo motor that can lock the running wheel, when commanded by the Raspberry Pi, which records wheel movement sensed by a rotary encoder mounted at the wheel shaft.

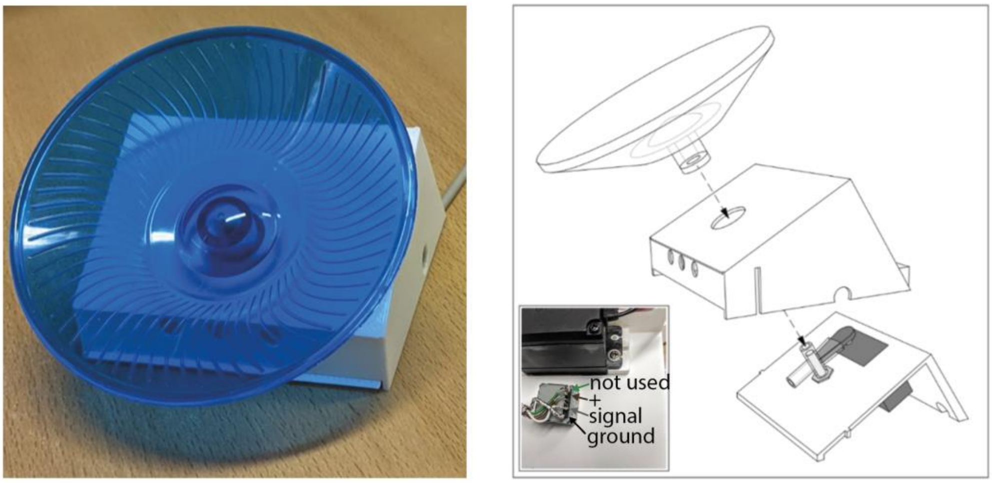


*Figure 19, Picture of the timed running wheel (left) and assembly geometry (right).*

A plastic 130 mm diameter wheel is removed from a commercially available (e.g., 'flying saucer') rodent running wheel. The shaft of a rotary encoder (Nidec Copal Electronics REC16B50-201) is fitted with a necessary amount of rubber tubing (typically a 20mm long piece of 5mm ID 7mm OD tubing is sufficient) to achieve a tight push-fit on the wheel's axis. The rotary encoder without the wheel is mounted in the 3-D printed enclosure base (rwA) along with the 180 deg servo motor (Master servo DS6020 C1689) which acts as the break (see Figure 19 for mounting geometry). A mounting nut and screws are supplied with the components. The supplied short hub of the servo is used, fitted with a soft piece of rubber or foam (e.g., a 20 mm piece of rubber tubing) to dampen contact with the wheel's axis. At this point the wiring should be completed in order to check functionality (see Electronics). Function and correct angle of the servo arm should be checked as follows: The movement arc of the servo is typically correct with the settings in the sample Arduino code, such that a rubber/foam attachment on the servo arm gently but firmly bends against the axis of the wheel, stopping its rotation. If this is not the case, the angle of the servo in the break and open position must be changed empirically by manually re-installing the servo arm at a different angle. After checking that the locking mechanism works normally, the servo and rotary encoder are covered with the second level of the enclosure (rwB). Lastly the wheel is fitted in place by pushing it onto the encoder's axis, making sure that the wheel can move freely and there is no friction between the rubber tubing and the enclosure. Superglue can be used to adhere the two parts (rwA and rwB) of the enclosure to each other. A 10 mm hole is drilled in the floor plate for the RW cable above its location (Figure 18A).

#### Electronics

See Figure 20 for connections. A Raspberry Pi 4b or 400 is connected to the load cell amplifier (SEN-13261) and RFID detector (SEN-09963) via USB cables (USB3.0 ports) and to the Arduino Mega through 12 digital lines. Additionally, the Arduino connects to the capacitive lick detector (input), visual output of detected licks (LED output, line D32), the servo motors (output) and the beam break sensors (input). The Raspberry Pi controls the Arduino and, directly, pellet dispensing from the FED3 (output) and water dispensing (output). Additionally the Raspberry Pi receives beam break and lick data from the Arduino, and, directly, pellet retrieval from the FED3 (input) and wheel rotation data (input).

Two-core shielded cable is used to connect servo motors, beam break detectors and FED3, and 4-core shielded cable for the timed running wheel and dispensing lick sensor. A switch (Adafruit P3064) is necessary on the servo power source, as the motors can damage components if powered up during start-up. This also makes troubleshooting easier as the motors can rapidly be turned off when needed and levers returned manually to safe operating range.

The 650nm Laser Diodes (SparkFun P1054) are connected to a separate 5V DC source (Adafruit P3064) as are beams and beam break sensors, which connect to the Arduino across 10 kOhm resistors. Importantly, all grounds should be wired together, apart from the 24V solenoid valve power source which is on a separate circuit.

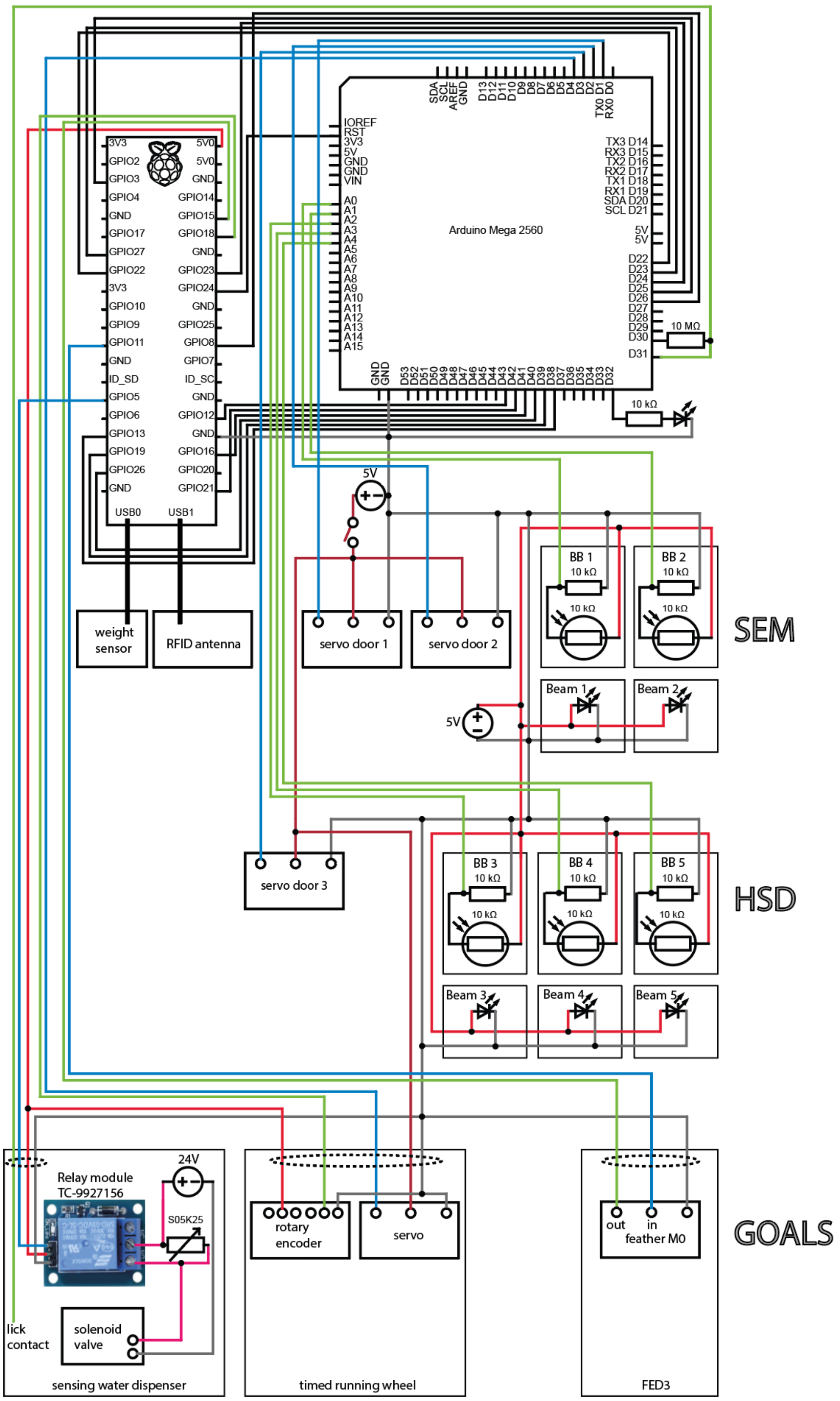


*Figure 20, Wiring diagram. Red = positive voltage source, green = sensor data, blue = output commands.*

**Code and software set up**

An Arduino Mega is set up for moving the servos, sensing the beam detectors and capacitive sensing of the lick spout. A Raspberry Pi is set up to control the experiment and log data. One can set up the software and code following these steps:

**1.** On the Raspberry Pi (we have used models 400 and 4b) with the operating system installed (Raspberry Pi OS, released 21-02-2023), Arduino IDE (version 1.8.19) is used to install the CapacitiveSensor[5] and Servo[6] libraries. This is done through the library installation menu under Tools>Manage Libraries… in Arduino IDE.

**2.** The Arduino code *Switch_maze_arduino_mega.ino* is downloaded from the supporting files or[7] and flashed to the Arduino Mega.

**3.** Normal operation of beam break sensors is tested empirically by uncommenting lines 173-182 (or sequentially each pair of lines to follow one BB at a time) in the Arduino code and flashing the code to the board. Then, enabling the serial monitor in Arduino IDE, values can be read out while occluding the sensor to simulate animal detection. The values for an open beam should be approximately the same in darkness and with room lights on. If this is not the case, the 3D-printed part holding the sensors can be painted black with, e.g., black nail polish, or a piece of transparent red plastic can be placed in front of the detector. The value for an occluded beam should be less than half the open beam value. If this is not the case, beam targeting may be improved while monitoring the values. Beam detection thresholds are set on lines 186, 195, 204, 213 and 222. After testing, lines 173-182 need to be commented out again to operate doors rapidly, and code flashed onto the board. For further adjustments and troubleshooting see below.

**4.** For setting up Python (Python 3.9.2 is preinstalled on the Raspberry Pi OS), Pandas and Numpy libraries are first installed via the terminal (e.g. `sudo pip3 install numpy` and `sudo pip3 install pandas`). The *requirements.pip* file is downloaded from[7] or supporting files of this document. After navigating in terminal to the folder containing the *requirements.pip* file, all listed requirements are installed by typing `pip3 install -r requirements.pip`.

**5.** All **.py* scripts and helper scripts should be downloaded from the supporting files or[7] and placed in the same folder.

**6.** On first installation, the scale needs to be calibrated and the correct settings must be set according to the manufacturer's instructions[9].
Briefly, after ensuring the correct USB port selection, a serial connection to the OpenScale is launched in Arduino IDE and the control menu is accessed by pressing 'x'. Baudrate is set to 9600, report rate to 120 ms, units to kg, decimals to 4, average amount to 1 and the serial trigger is switched 'off'. The scale is then tared to zero and calibrated with a 25 g weight. Calibration should be repeated daily in the beginning as the load cell can 'creep' somewhat after installation.

**7.** The angles that a servo needs to achieve to close and open a door will likely vary based on the exact distances used in each build, and are set by trial and error on lines 35-40 of the Arduino code. As a standard practice, install the levers such that they cannot hit the floor board above them, i.e., power off the servo, turn the servo hub to the extreme position and install the lever there such that it is near but not touching the floor. Because we do this, the default code has 'closed' values of 179 degrees. After that it will be simple to find a suitable 'open' value by testing values less than 90 degrees. In case the door lever collides with something, turn the power to the servos off rapidly from their power switch, go back to the code and bring the degree value closer to 90. When powering up the system the servos can receive extreme commands. Therefore it is best to power up the servos last.

**8.** Two manual setup operations must be done every time after restarting the Pi:

1. The PiGPIO daemon is launched from the terminal (`sudo pigpiod`).
2. The servos are powered up and door angles are checked/changed for safe operation (see section above).

**9.** Now the main script can be run via the terminal or an IDE (we use Thonny Python IDE).

**Adjustments and troubleshooting**

***Beam targeting***

Beam break devices in the SEM are targeted as explained in [1], briefly: Beam break 1 is used for safely closing door 1: Beam 1 is targeted above door 1 such that when the door is open the beam hits the photoresistor. When the door is closed or an animal is on top of the open door, the beam is broken. Beam break 2 (BB2) is used for detecting an exiting animal at a safe distance from door 2 (>100 mm): Beam 2 is targeted through the sides of the single entry module such that an animal in the middle of the module will break it.

Beam break devices in the HSD (BB3-5) are used like BB2 to detect an animal when the beam is broken. These enable safe operation of the HSD and logging entries to the goals or start point. These beams should be at a height of approximately 15 mm above the floor.

Beam detector readings can be monitored in Arduino IDE serial monitor after uncommenting lines 173-182 in the Arduino code (or to see only the initial calibration readings, lines 146-155). After diagnostics, these lines need to be commented out again to operate doors rapidly. Beam detection thresholds are set on lines 186, 195, 204, 213 and 222.

***Weight sensor parameter selection and calibration***

The weight sensor is set up and calibrated according to the manufacturer's instructions[9]. Briefly, after ensuring the correct USB port selection, a serial connection to the OpenScale is launched in Arduino IDE and the control menu is accessed by pressing 'x'. Baudrate is set to 9600, report rate to 120 ms, units to kg, decimals to 4, average amount to 1 and the serial trigger is switched 'off'. The scale is then tared to zero and calibrated with a 25 g weight. Calibration should be repeated daily in the beginning as the load cell can 'creep' somewhat after installation.

***Door speed***

The speed of door movements is set by the slowness constant on line 13 of the Arduino code. This value corresponds to the time, in microseconds, it takes to move the servo by 1 degree. We have found 150 ideal.

#### Using the Switchmaze

##### Radio-frequency identification (RFID) tag implantation

30 wild-type C57BL6 male mice were used in this study. All experimental procedures were approved by the Netherlands Central Committee for Animal Experiments and the Animal Ethical Care Committee of the Vrije Universiteit Amsterdam (AVD11200202114477). To implant a glass RFID tag capsule under the skin, each mouse was anesthetized with sleep mix (i.p., fentanyl 0.05 mg/kg, medetomidine 0.5 mg/kg and midazolam 5 mg/kg in saline). An RFID chip (Sparkfun SEN-09416) was implanted under the chest skin using a non-medical ID transponder syringe RFID injector (e.g., DHgate 533480816). The wound was closed with tissue glue, anaesthesia was antagonized with wake mix (i.p., flumazenil 0.1 mg/ml and atipamezole 5 mg/ml in saline) and the animal received 0.05 mg/ml carprofen in drinking water for 2-4 days as post-operative pain medication.

##### Reading RFID tags for the first time and habituating animals

To test the Switchmaze, mice were housed in the apparatus for up to a month in cohorts of 2-4 animals. RFID tagged animals were first read in by allowing animals to explore the open SEM while running the script *RFIDreader_newcohort_main.py*. After starting the script, RFID tagged animals were placed in the Switchmaze in their home cage and the nest wall assembly was put in place. After the RFID tags of all animals were detected (10 - 60 min of free exploration typically), the tags were written into lines 27-30 of *Switch_maze_functions.py* and normal operation was started by running *Switch_maze_main.py*. For the first two days, water was available *ad libitum* in the home cage and 2g/animal of dry chow was provided on the home cage floor. This was done to ensure adequate food and water intake during the time when entering the foraging environment is a new action. Therefore, mice would first enter the foraging environment due to exploration. In 6-48 h, they learned to use the goal objects and obtain food and water from the maze.

During the initial habituation period, there were very infrequent events where an animal got stuck in the foraging environment after a maze exit was triggered. This was likely due to the animal making an elongated probing posture while gripping a closing door with its hindleg. These situations were noticed by monitoring via an overhead camera and the incoming event data, and resolved by running the script *rescue_main.py* which stops maze operation, waits for a complete exit and reports it via email to the user. We have also included a version of the routine operation code that sends an email warning to the user if an animal's body weight drops below 85% of a user-defined reference weight. As this may be less stable than the standard code, we advise testing it after achieving routine operation. This code can be found in the repository as *Switch_maze_functions_email.py* and *Switch_maze_main_email.py*.

During baseline operation, an animal typically consumed 189 ± 49 food pellets (14 mg) in a 24 h period, which corresponds to the typical number of small meals wild mice are estimated to eat in a night[10]. Upon most entries into the food pod, a pellet was consumed (93.9 ± 10.1 % of entries). 210 ± 130 water drops (10 µl) were retrieved and drinking occurred upon 98.5 ± 2.3 % of entries into the water pod. In a 24 h period, an animal ran on the running wheel on average 6.7 ± 13.3 revolutions during 5 ± 7 entries into the water pod which accounted for 4.0 ± 6.5 % of the entries. Therefore, despite the appeal of running wheels even to wild mice[11], use of the running wheel was marginal and is unlikely to affect the results. Food (201 ± 49) and water (212 ± 130) pod entries were approximately balanced over a 24 h period.

### Histology

Below are the histology figures showing all H-hM4Di and PFC-hM4Di animals, in addition to three example control animals.

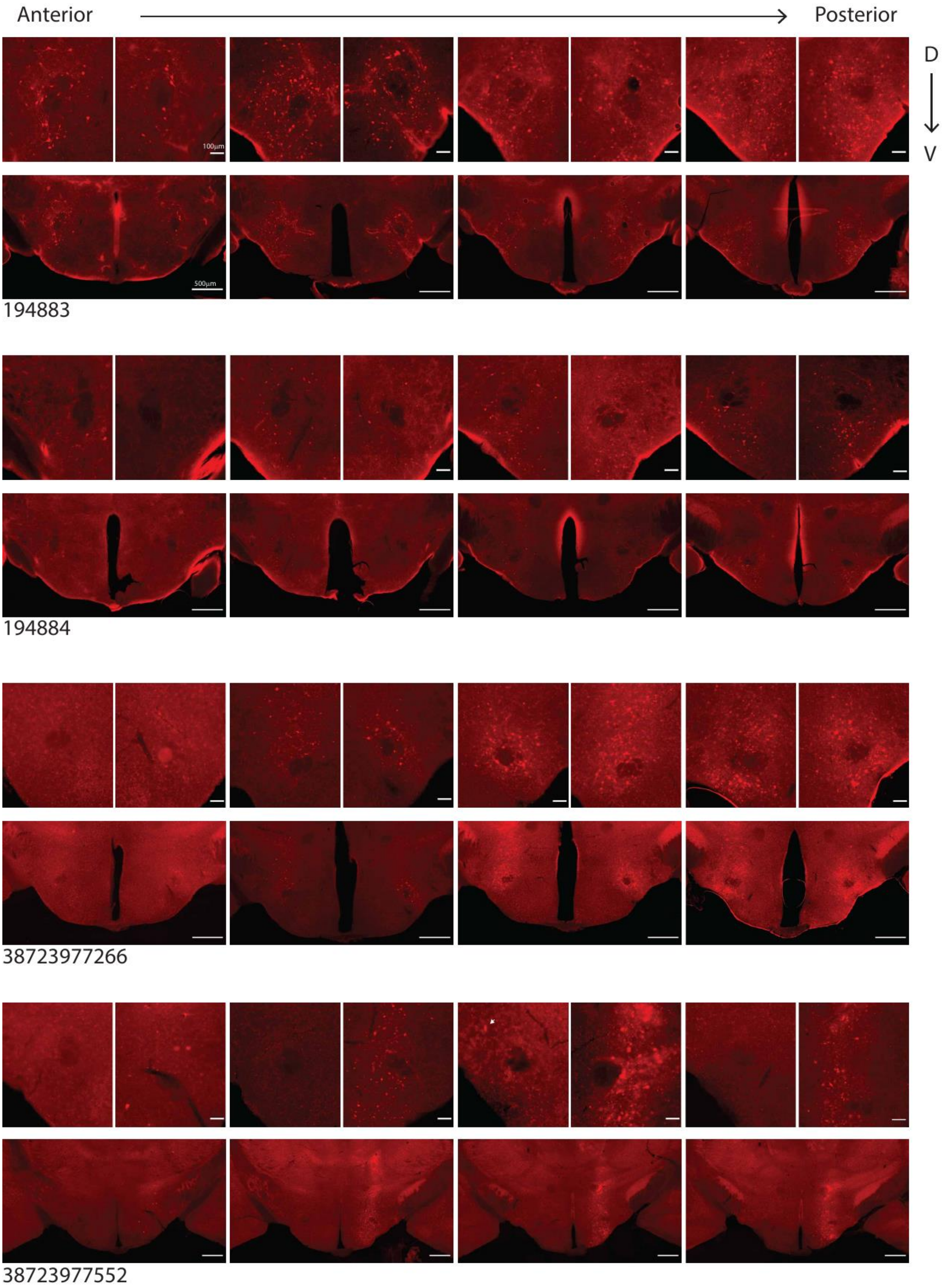


Figure 1, Expression of red fluorescent protein in H-hM4Di cases and lack of expression in two controls. For each animal, two levels of magnification are provided (top row, high magnification, and bottom row low) showing the lateral hypothalamus. Arrows point to difficult-to-detect expression.

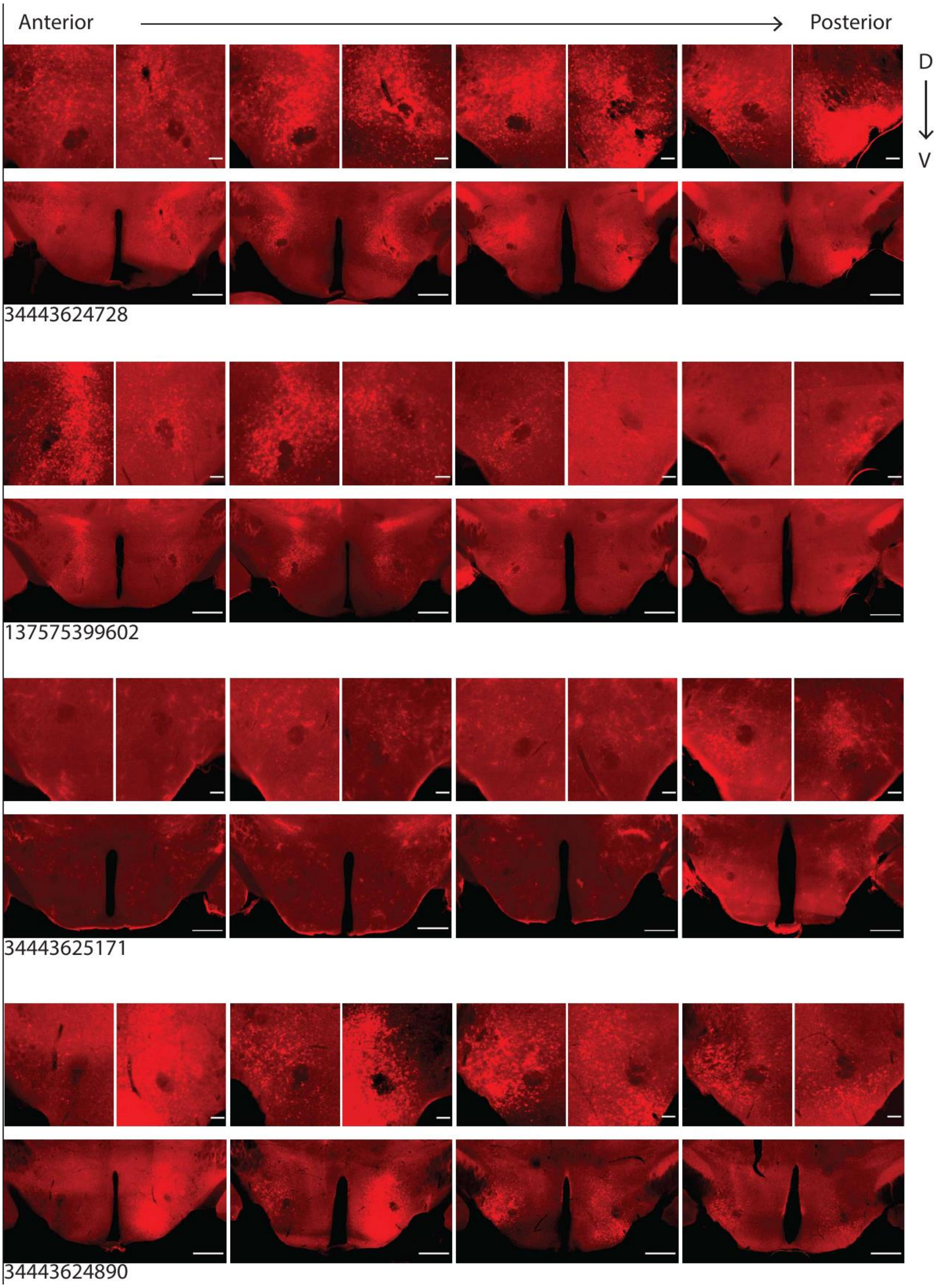


Figure 1, continued.


bioRxiv THE PREPRINT SERVER FOR BIOLOGY

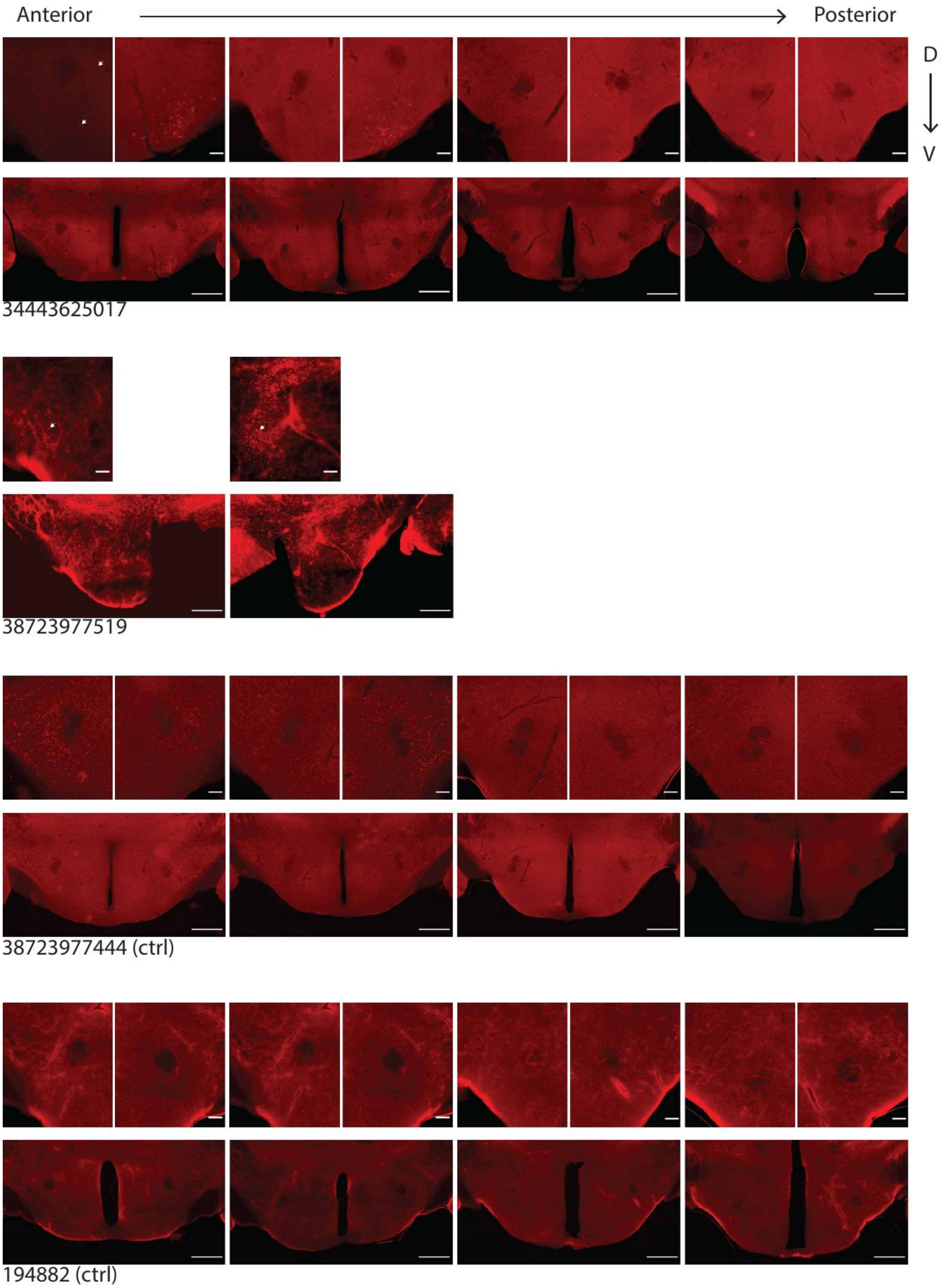


Figure 1, continued.

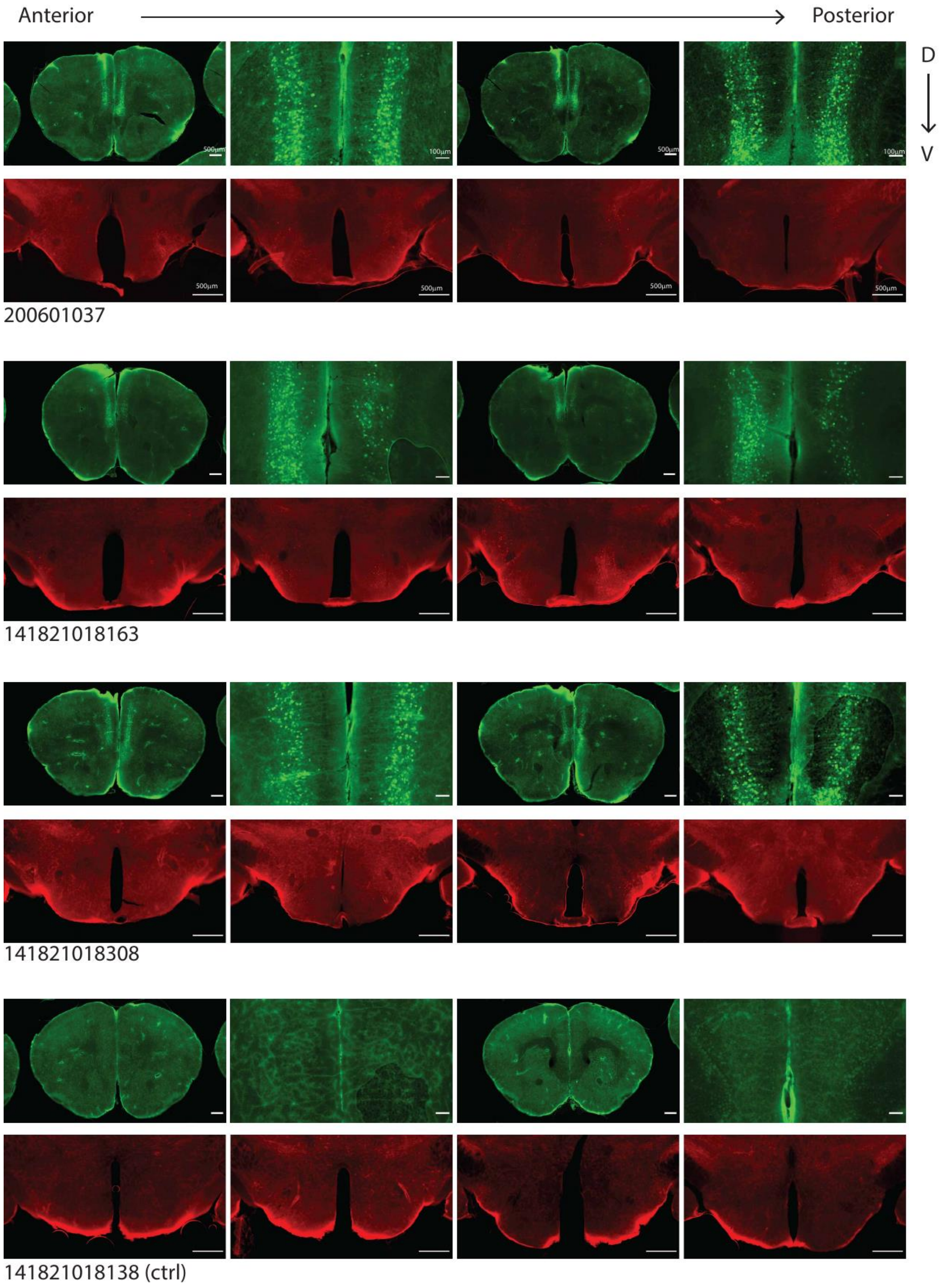


Figure 2, Expression of tdTomato and mCitrine in all PFC-hM4Di cases and lack of expression in four controls. For each animal, two levels of magnification are provided for PFC (top row left to right: two pairs of low, then high magnification images, and bottom row low), and one level of magnification for the LH (bottom rows). Arrows point to difficult-to-detect expression.

 bioRxiv THE PREPRINT SERVER FOR BIOLOGY 

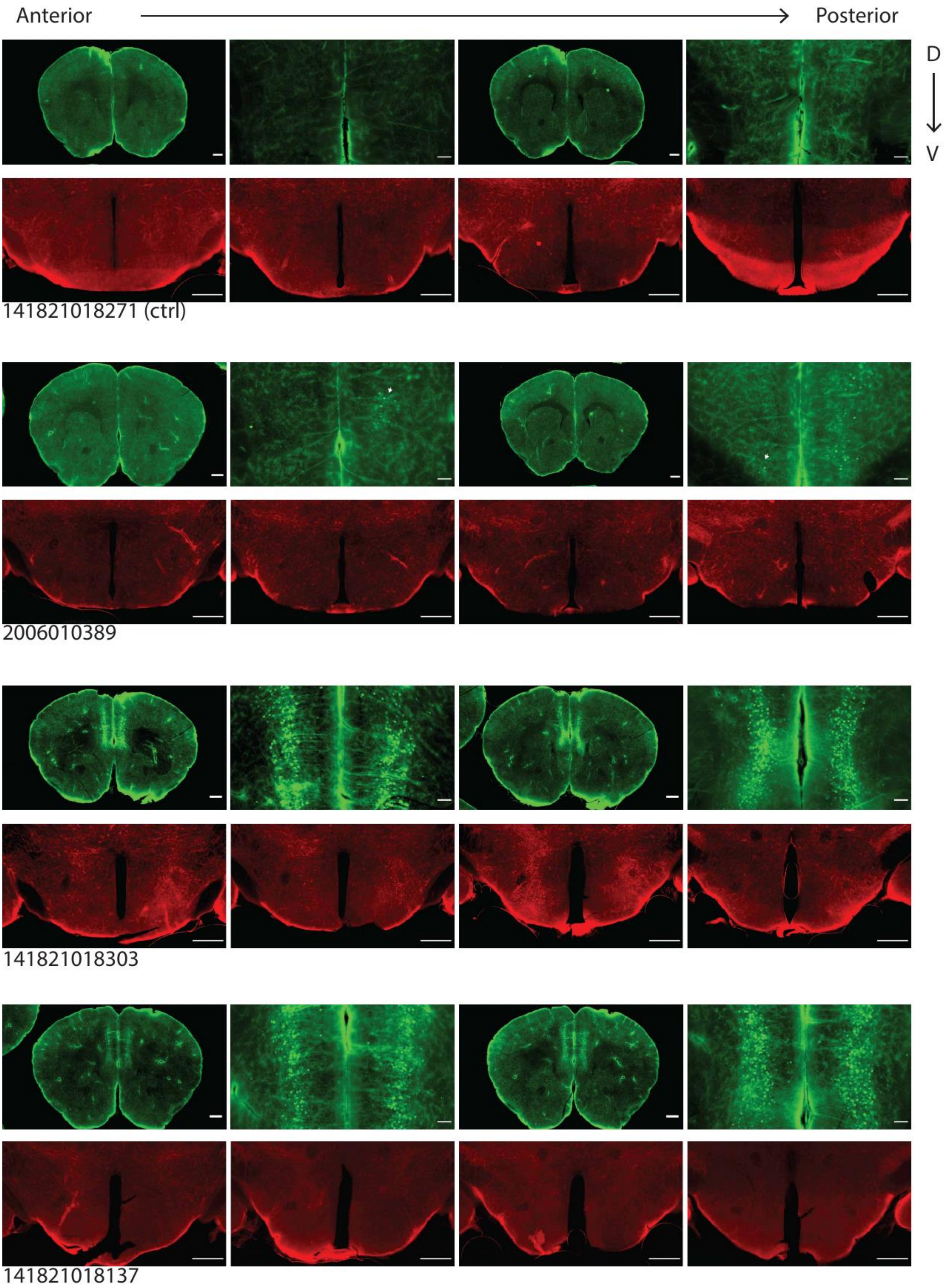


Figure 2, continued.

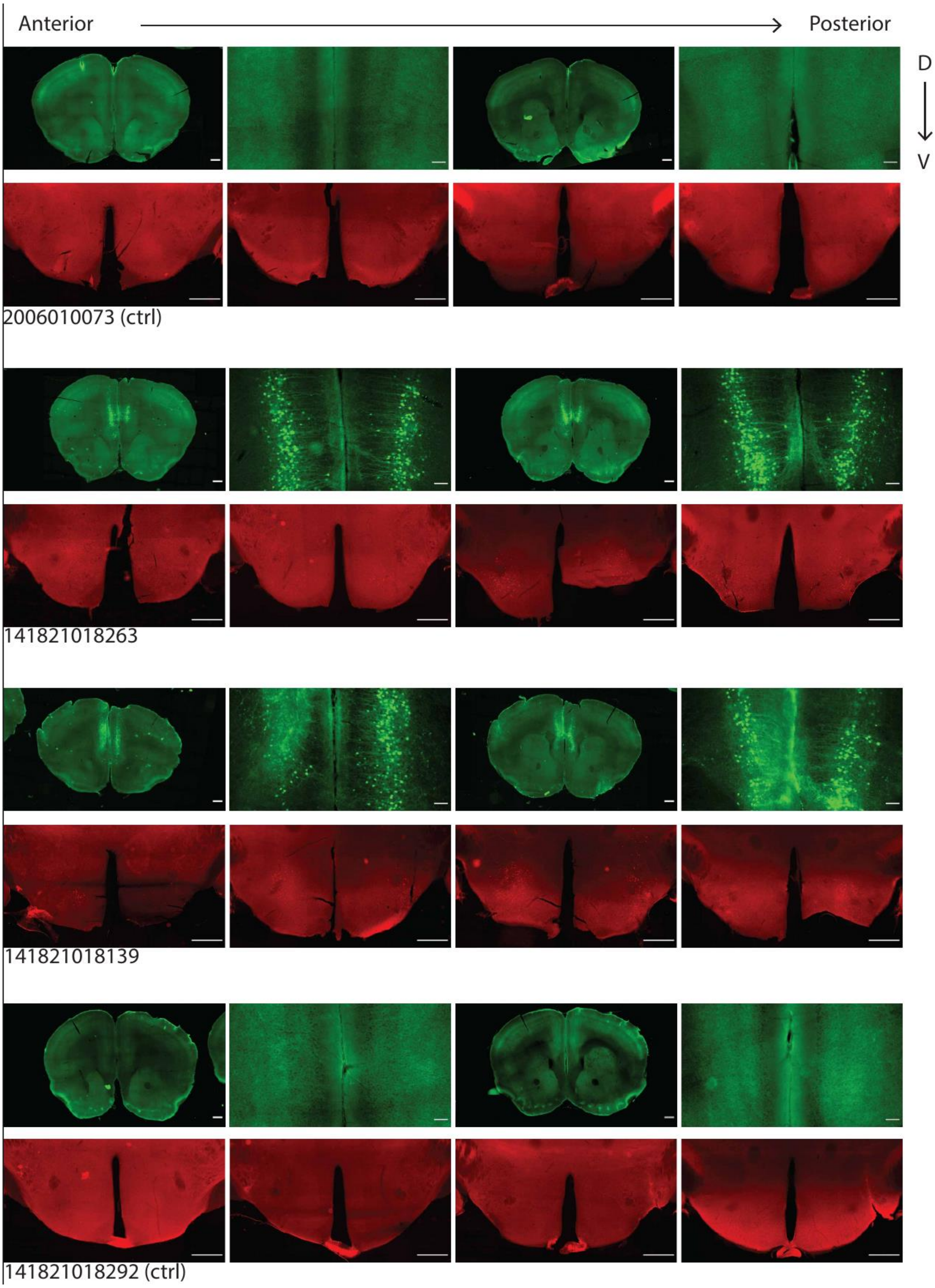


Figure 2, continued.

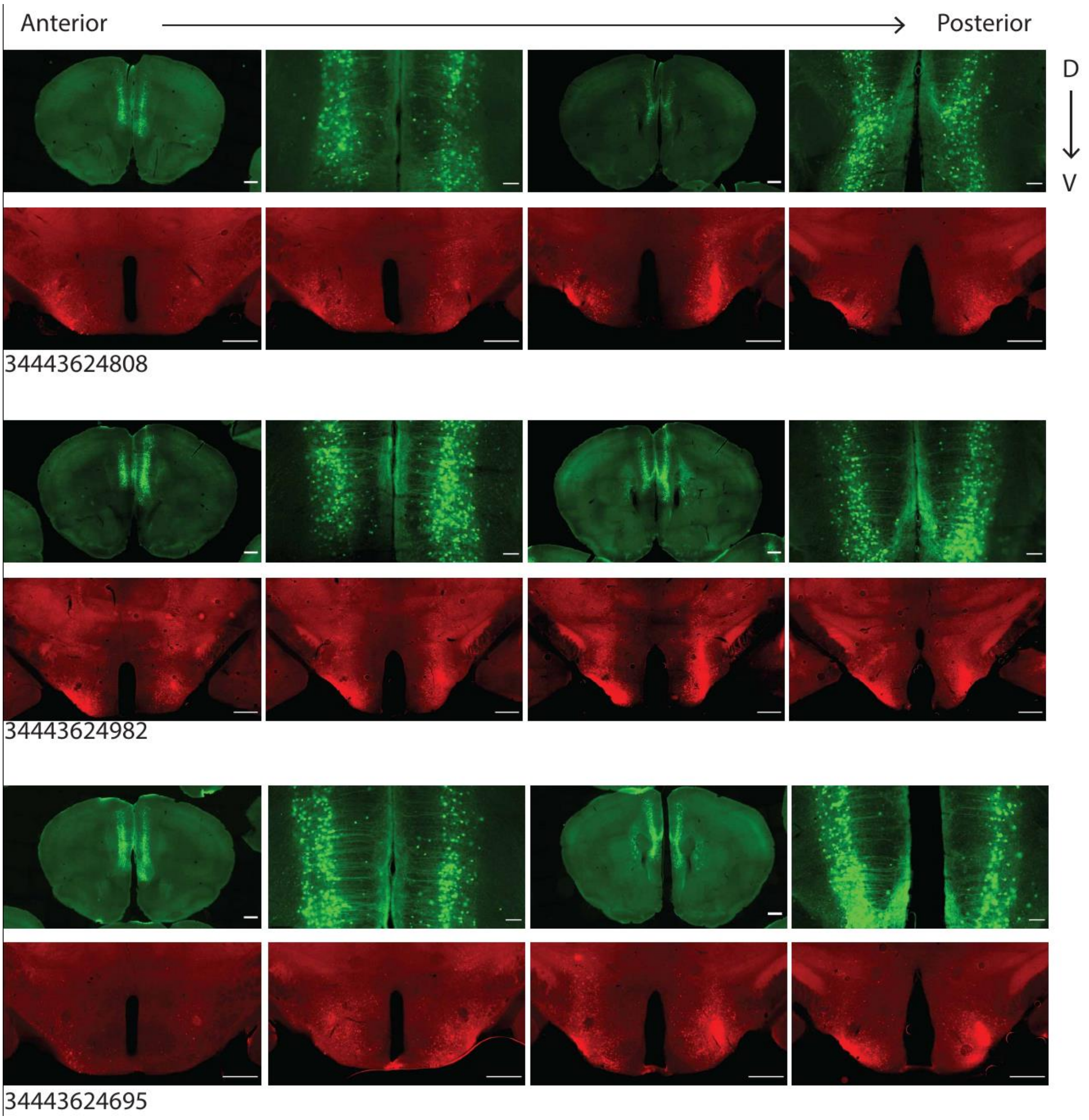


Figure 2, continued.
